# Stress conditions promote the mating competency of *Leishmania* promastigotes *in vitro* marked by expression of the ancestral gamete fusogen HAP2

**DOI:** 10.1101/2021.08.31.458317

**Authors:** Isabelle Louradour, Tiago Rodrigues Ferreira, Emma Duge, Nadira Karunaweera, Andrea Paun, David Sacks

**Author notes:** Correspondence should be addressed to D. S. these authors contributed equally (Isabelle Louradour, Tiago Rodrigues Ferreira). Department of Parasites and Insect Vectors, Institut Pasteur/INSERM U1201, 75015 Paris, France (Isabelle Louradour).

## Abstract

*Leishmania* are protozoan parasites transmitted by the bite of sand fly vectors producing a wide spectrum of diseases in their mammalian hosts. These diverse clinical outcomes are directly associated with parasite strain and species diversity. Although *Leishmania* reproduction is mainly clonal, a cryptic sexual cycle capable of producing hybrid genotypes has been inferred from population genetic studies, and directly demonstrated by laboratory crosses. Experimentally, mating competency has been largely confined to promastigotes developing in the sand fly midgut. The ability to hybridize culture promastigotes *in vitro* has been limited so far to low efficiency mating between two *L. tropica* strains, L747 and MA37, that mate with high efficiency in flies. Here, we show that exposure of promastigote cultures to DNA damage stress produces a remarkably enhanced efficiency of *in vitro* hybridization of the *L. tropica* strains, and extends to other species, including *L. donovani, L. infantum*, and *L. braziliensis*, a capacity to generate intra- and interspecific hybrids. Whole genome sequencing and total DNA content analyses indicate that the hybrids are in each case full genome, polyploid hybrids. Single-cell RNA sequencing of the L747 and MA37 parental lines highlights the transcriptome heterogeneity of culture promastigotes and reveals discrete clusters that emerge post-irradiation in which genes potentially involved in genetic exchange are expressed, including the ancestral gamete fusogen *HAP2*. By generating reporter constructs for HAP2, we could select for mating-competent and mating-incompetent promastigotes. Overall, this work reveals that there are specific populations involved in *Leishmania* mating associated with a discernible transcriptomic signature, and that stress facilitated *in vitro* hybridization can be a transformative approach to generate large numbers of hybrid genotypes between diverse species and strains.

## Introduction

Protozoan parasites of the *Leishmania* genus produce a spectrum of diseases in their mammalian hosts, including humans, ranging from self-limiting cutaneous lesions to tissue destructive mucosal involvement to infection of the deep viscera that is fatal in the absence of treatment. These clinical outcomes are associated with an extraordinary diversity of *Leishmania* species and strains, with more than 20 species that are pathogenic to humans. All members of the genus have a dimorphic life cycle consisting of extracellular promastigotes that multiply asexually in the digestive tract of their sand fly vectors, and amastigotes that multiply asexually in the phagosomes of their mammalian host cells, primarily macrophages. *Leishmania* were long considered to be essentially clonal, with genetic diversity arising from gradual accumulation of somatic mutations (1).

The reproductive strategies of *Leishmania* are now known to include a cryptic sexual cycle that has been inferred from the analysis of hybrid genotypes observed in natural isolates (2-7), and directly demonstrated by the generation of hybrids between different *Leishmania* strains and species in the laboratory (8-11). The later involved mating between extracellular promastigote stages developing in the sand fly, and was achieved by co-infection of flies with two parental lines bearing different drug-resistance markers, with subsequent selection of double-drug resistant cells. Experimentally at least, the sexual cycle is non-obligatory, relatively rare, and confined to life-cycle stages present in the sand fly midgut. Based on whole genome sequencing analyses, the allele inheritance patterns of experimental hybrids provides strong evidence that the system of genetic exchange in *Leishmania* is Mendelian and involves meiosis-like sexual recombination (12). Homologues of meiosis-specific genes are found within the genomes of *Leishmania*, and are expressed by stages in the sand fly, including the core meiotic genes *spo11, hop1* and *dmc1* involved in creating DNA double-strand breaks, homologous chromosome alignment and recombination (13, 14). Homologues of the cell and nuclear fusion proteins HAP2 and GEX1, respectively, have also been found in *Leishmania* ancestral lineages, providing indirect evidence that sex was already present in the last eukaryotic common ancestor (15). Importantly, there has been no direct evidence that any of these genes or their products are involved in *Leishmania* genetic exchange, and due to the difficulty of making direct observations of the mating events in the sand fly, the precise nature of this reproductive process, including the identification of putative gametic cells, has been difficult to study.

We recently reported that *Leishmania* promastigotes, which can be readily grown in axenic culture, can form stable hybrids entirely in vitro (16). While this result proved that mating competent cells can arise in culture, its broader applicability to study the biology of mating and to generate recombinant parasites for genetic linkage analysis was limited by the fact that the *in vitro* hybridization was confined to only two strains of *L. tropica*, and occurred at frequencies far lower than in the sand fly. In the current studies, we have investigated the conditions that might potentiate the mating success of *Leishmania in vitro*. It is common for organisms that are facultatively sexual to undergo sex in response to environmental stress, including conditions that produce DNA damage (17, 18). Thus, a primary function of sex is proposed as an adaptation for DNA repair, for which a homologous, undamaged chromosome serves as a template (19). Amongst eukaryotic microbes, oxidative DNA damaging conditions have been shown to promote sexual spore formation in the yeast *Schizosaccharomyces pombe* (20) and in the multicellular green alga *Volvox carteri* (21). Oxidative stress also efficiently induced the expression of sexual pheromone precursors and same-sex mating in *Candida albicans* (22). DNA damage caused by X-irradiation induced meiotic recombination in the budding yeast *Saccharomyces cerevisiae* (23) and in the nematode *Caenorhabditis elegans* (24). In the current studies, we have used DNA damaging conditions to promote *in vitro* mating of *Leishmania*. Exposure to H_2_O_2,_ methyl methanesulfonate (MMS), or to γ-radiation produced a remarkably enhanced efficiency in the *in vitro* hybridization of the *L. tropica* strains, and extended to other species and strains a capacity for *in vitro* mating. Single cell RNA-seq comparisons of untreated and irradiated promastigotes revealed greatly expanded subpopulations amongst the irradiated cells that expressed meiotic gene homologues. We identified expression of *HAP2* as the main meiotic gene associated with enhanced mating that could be used to identify and select for mating-competent cells in culture.

## Results

### Stress conditions increase the frequency of *Leishmania* hybrid formation *in vitro*

We recently developed a protocol for generating *Leishmania* hybrids completely *in vitro* using two *L*.*tropica* strains, L747 and MA37, each transfected with different drug resistance and fluorescent markers (16). While this protocol achieved the recovery of stable, full genome hybrids, referred to as LMA hybrids, the frequency of *in vitro* hybridization was notably low, particularly in comparison to the frequency of LMA hybrids generated in sand fly co-infection experiments. Stress conditions leading to DNA damage have been shown in other organisms to trigger their sexual reproductive cycles (17-19). Therefore, we chose exposure to γ-radiation, hydrogen peroxide (H_2_O_2_), or methyl methanesulfonate (MMS), treatments known for their genotoxic effects, as experimental stress conditions. To assess the effects of these treatments on the frequency of hybrid formation *in vitro*, both the L747 RFP-Hyg and MA37 GFP-Neo parental lines were pretreated or not with the various conditions, and were then were mixed in equal volumes without any drug selection and distributed into 96-well plates. After 24 hrs, the content of each well was transferred into double-drug selective medium (Hygromycin and Neomycin) in 24-well plates. In untreated conditions, a low proportion (1.7%) of wells yielded double-drug resistant lines (Fig. 1A-C). When L747 and MA37 parental lines were pre-exposed to either γ-radiation, H_2_O_2_ or MMS, the frequency of positive wells averaged 63.7%, 26.5%, or 34.5%, respectively, in the 4 independent experiments conducted for each treatment. These increased frequencies were observed despite the fact that fewer of the pre-treated L747 and MA37 cells were present at the initiation of co-culture, due to the negative impact of each of these treatment on parasite growth. Growth curves of L747 and MA37 parasites cultures after exposure to γ-radiation, H_2_O_2_ or MMS are shown (Sup. Fig. 1). Area under the curve (AUC) analysis of the culture growth showed a total area ratio of 0.71 fold and 0.44 fold in irradiated L747 and MA37 cultures, respectively, compared to their respective untreated controls. The minimal frequency of mating competent cells for the two parental lines, calculated using the number of input cells at the start of each co-culture and assuming that each positive well contained only one hybridization event, was between 10^−8^ and 10^− 9^ in untreated conditions (Fig. 1 D-F; Sup. Table 1). These frequencies increased between 59 and 347 fold when the cultures were submitted to the different stress conditions.

**Figure 1.**
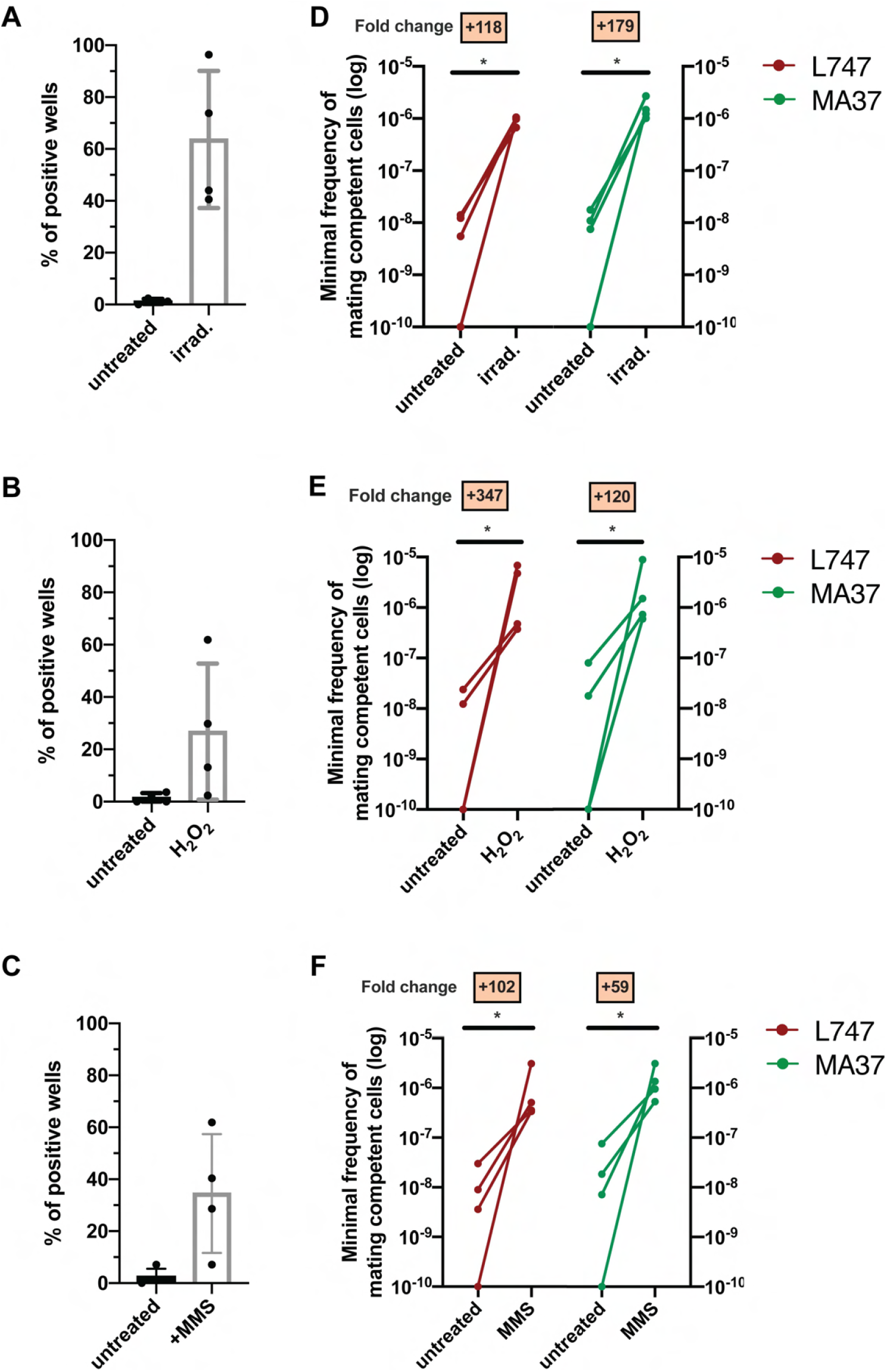
Stress treatments strongly enhance *in vitro* mating of *L. tropica* L747 and MA37. (**A-C**) Proportion of culture wells exhibiting growth of LMA hybrids when exposed to γ-radiation (A), hydrogen peroxide (H_2_O_2_) (B), or methyl methanesulfonate (MMS) (C). (**D-E**): Minimum frequency of mating competent cells for both L747 and MA37 parental strains after exposure to γ-radiation (D), H_2_O_2_ (E) or MMS (F). Four independent experiments were performed for each treatment. The frequencies calculated for the control and treated conditions in each experiment are linked by a line. \**p* = 0.0286 (Mann-Whitney test).

### Pre-exposure to γ-radiation promotes the generation of intraspecific and interspecific *Leishmania* hybrids

Our previous attempts to generate *in vitro* hybrids involving *Leishmania* strains other than *L*.*tropica* L747 and MA37 were unsuccessful (16). To evaluate the applicability of the current protocol to different species, we performed a series of crosses using different pairs of parental lines exposed or not to γ-radiation. We attempted several intraspecific crosses: *L. tropica* (*Lt*) MA37 x *Lt*Moro; *L. donovani* (*Ld*) Mongi x *Ld*SL2706; *L. braziliensis* (*Lb*) RicX x *Lb*M1; and *L. major* (*Lmj*) FV1 x *Lmj*LV39 (Table 1). Doubly-drug resistant lines were successfully generated from all of these mating pairs, with the exception of *Lmj*FV1 and *Lmj*LV39. In addition, cross-species hybrids were recovered from *Lt*MA37 and *L. infantum* (*Li*) LLM320 (Table 1). All of the hybrids generated in these crosses were obtained using irradiated parents, with the exception of a single *Lt*MA37 x *Lt*Moro hybrid which was generated using untreated cells. As the parental lines also expressed GFP or RFP fluorescence markers, we used flow cytometry as a rapid way to confirm the hybrid nature of the parasites growing in our assays prior to cloning (Sup. Fig. 2). We also compared irradiated *L. braziliensis* crosses in a construct-swap approach. Comparatively, the frequencies of hybrid recovery for both MIR (*Lb*M1-RFPHyg x *Lb*RicX-GFPNeo) and RIM (*Lb*RicX-RFPHyg x *Lb*M1-GFPNeo) crosses were identical (Table 1), as well as the overall parental allele contribution (Sup. Fig. 3), suggesting the reproducibility of this protocol independently of the pairwise combination of the two drug resistance markers in the two parental strains.

**Table 1.** List of the different intra- and interspecific irradiation-facilitated hybrids (from *in vitro* mating after exposure of parental strains to gamma-radiation) generated in this study with their respective ploidies according to propidium iodide staining, flow cytometry and whole-genome sequencing.

| Type of cross | Parental strains |  | Hybrid denomination | Frequency of hybrid recovery: # positive wells / # total wells |  | Ploidy of irradiation-facilitated hybrids: # hybrids / # hybrids tested |  |  |
| --- | --- | --- | --- | --- | --- | --- | --- | --- |
|  | A | B |  | No treatment | Irradiation | 2n | 3n | 4n |
| intraspecific | *L.tropica* L747 RFP-Hyg | *L.tropica* MA37 GFP-Neo | LMA | 4/336 | 214/336 | 0/44 | 4/44 | 40/44 |
| intraspecific | *L.tropica* Moro RFP-Hyg | *L.tropica* MA37 GFP-Neo | MoMA | 1/84 | 20/84 | 0/4 | 1/4 | 3/4 |
| interspecific | *L.infantum* LLM320 RFP-Hyg | *L.tropica* MA37 GFP-Neo | IMA | 0/84 | 3/84 | 0/3 | 0/3 | 3/3 |
| intraspecific | *L.donovani* Mongi RFP-Hyg | *L.donovani* SL2706 GFP-Neo | Mondo06 | 0/84 | 35/84 | 0/16 | 0/16 | 16/16 |
| intraspecific | *L.braziliensis* M1 RFP-Hyg | *L.braziliensis* RicX GFP-Neo | MIR | 0/84 | 9/84 | 0/7 | 1/7 | 6/7 |
| intraspecific | *L.braziliensis* RicX RFP-Hyg | *L.braziliensis* M1 GFP-Neo | RIM | 0/84 | 9/84 | 0/6 | 0/6 | 6/6 |
| intraspecific | *L.major* FV1 Sat | *L.major* LV39 RFP-Hyg | LVFV | 0/84 | 0/84 | * | * | * |

RFP: red fluorescent protein; GFP: green fluorescent protein; mNG-yTub: mNeonGreen-gammaTubulin fusion; Hyg: hygromycin B resistance marker; Neo: neomycin resistance marker; Bsd: blasticind S resistance marker; N.D.: not detected

### Genome ploidy and whole genome sequencing analyses reveal that *in vitro* hybrids are full-genome, polyploid hybrids

The double-drug resistant hybrid lines were cloned by distribution in 96 well plates in promastigotes growth medium containing both antibiotics. The DNA content of selected irradiation-facilitated, *in vitro* hybrid clones was analyzed using propidium iodide (PI) staining followed by flow cytometry. The diploid parental lines were used as controls for normalization. With the exception of 4 triploid LMA hybrids, 1 triploid *Lt*MA37 x *Lt*Moro (MoMA) hybrid and 1 triploid *L. braziliensis* hybrid, all of the other intra- and interspecific hybrids were close to tetraploid (Fig. 2A, Sup. Fig. 4, Table 1). No diploid hybrids were observed, which distinguishes them from the LMA hybrids recovered from the *in vitro* LMA hybrids generated from untreated parental lines, for which diploid and polyploid hybrids were obtained (16). Confocal microscopy imaging of Hoechst 33342 stained promastigotes and transmission electron microscopy show that the irradiation-facilitated hybrids carry a single nucleus and are not heterokaryons (Fig. 2B,C).

**Figure 2.**
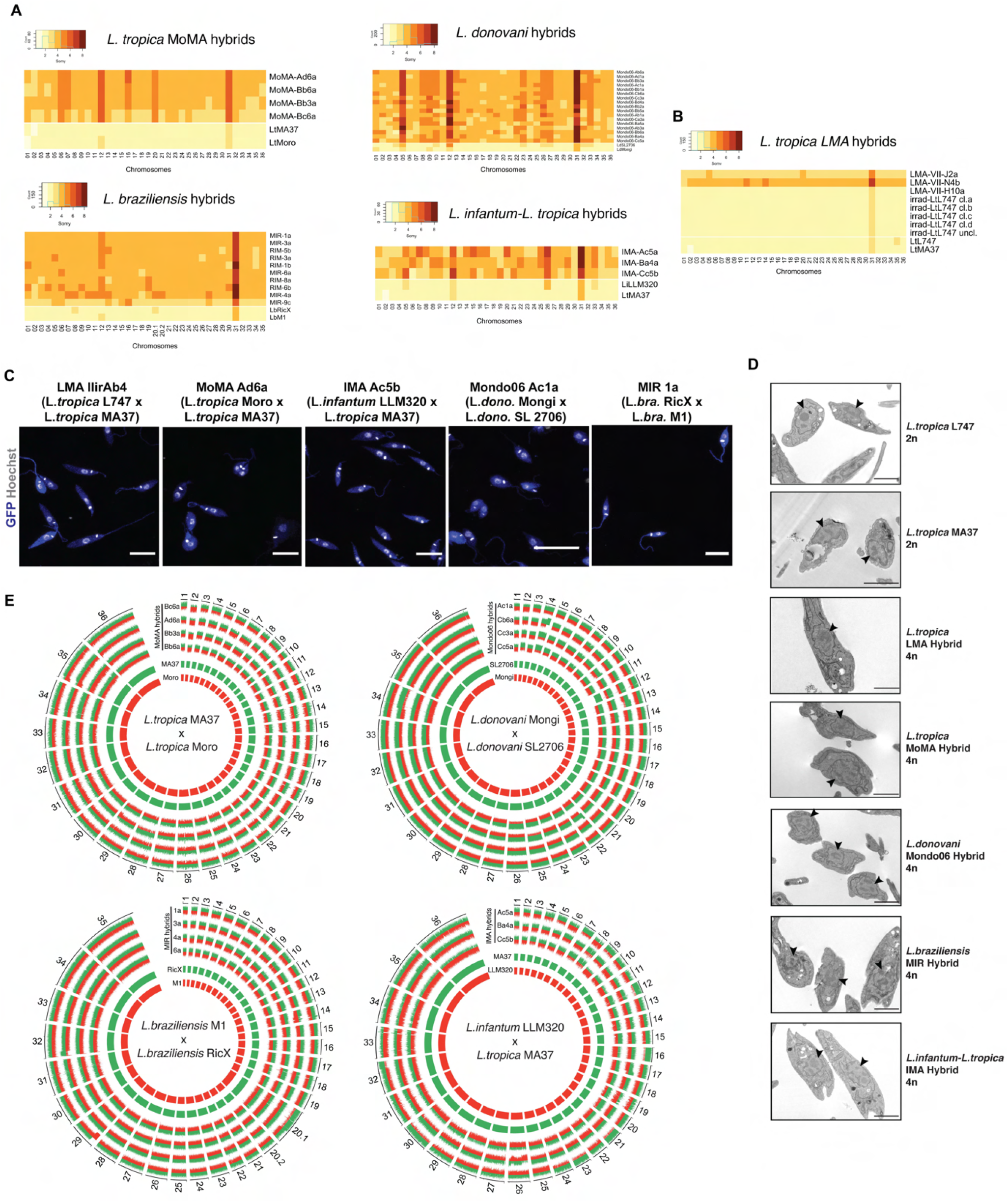
Whole-genome sequencing of irradiation-facilitated experimental hybrids reveals chromosome biparental inheritance and recombination breakpoints. (**A**) Normalized sequence read depth of the different *Leishmania* chromosomes (35 chrs in *L. braziliensis* and 36 in the other species shown) was used to infer somy values in parental strains (bottom two rows in each heatmap) and hybrids. **(B)** Normalized sequence read depth of *L. tropica* LMA hybrids of different ploidies generated in untreated *in vitro* mating, and of 4 independent sub-clones and one uncloned line of the irradiated L747 parent. (**C**) Fluorescence confocal microscopy images of tetraploid LMA, MoMA, IMA, Mondo06 and MIR irradiation-facilitated hybrids. Nuclear and mitochondrial DNA (kDNA) are shown in white (Hoechst 33342 staining) and cell bodies in blue (GFP expression). Scale bar: 10µm. (**D**) Transmission electron micrographs of representative diploid (2n) parental strains and different tetraploid (4n) hybrids depicting a single nucleus (arrowheads) in promastigote forms not undergoing cell division. Scale bar: 2 µm. (**E**) Circos plots representing the inheritance patterns of all the homozygous SNP differences between the two parental strains in four *L. tropica* MoMA intraspecific hybrids, three *L. infantum-L. tropica* IMA interspecific hybrids, four *L. donovani* Mondo06 intraspecific hybrids and in four *L. braziliensis* MIR intraspecific hybrids. Each group separated by radial white lines represents a different chromosome and chromosome ids are shown on the outer circle. Hybrid clones ids are indicated at the start of each circular track. Red and green histograms depict inferred parental contribution from homozygous SNPs specific to the RFP-expressing parental line and the GFP-expressing parental line, respectively. Whole-genome sequencing analyses were performed using the PAINT software (53) and reference genomes available on Tritrypdb (tritrypdb.org).

While polyploid hybrids, including tetraploid progeny, have been recovered from experimental hybrid lines generated in sand flies, the majority of *in vivo* hybrids have been close to diploid (8, 9, 12). These results suggest that tetraploid intermediates, as have been described in some fungi, might reflect a normal part of the sexual cycle in *Leishmania*. In this context, under the conditions of irradiation-facilitated hybridization *in vitro*, promastigotes capable of cell and nuclear fusion can be generated but somehow fail to complete the ploidy reduction steps of the meiotic cycle. In attempts to promote these processes, selected LMA tetraploid clones were re-exposed to stress conditions, including H_2_O_2_ and γ-radiation, and/or were passed through sand flies. All of these attempts to induce ploidy changes in the LMA hybrids were unsuccessful (Sup. Fig. 5).

To determine the copy number of individual chromosomes and their respective parental genome contributions, we performed whole genome sequencing of selected progeny clones. For somy quantification, the average of coverage for each chromosome was scaled to the ploidy of the cells. The somy profiles of the parents and hybrid clones are depicted as heat maps in Fig. 2A. The parental somies are in each case close to 2n, with the exception of chr 31 which is trisomic in every parent, along with other chromosomes that show close to 3n profiles in combinations that are specific to the parental line, e.g. chrs 8, 20.1 in *Lb*RicX; chrs 5, 12, 26, 33 in *Li*LLM320. The majority of the chromosomes that are 2n in both parents are close to 4n in the hybrid progeny. For those chromosomes that are close to 3n in one of the parents, the hybrid somies are in most cases close to 5n, while chromosomes that are close to 3n in both parents, are close to 6n in the hybrids. These inheritance patterns reinforce the conclusion that the hybrids are products of the fusion of the complete parental genomes. There are, nonetheless, a large number of hybrid chromosomes that deviate from the expected additive copy number, showing an apparent gain or loss of somy. As a result, within a particular cross each of the hybrids appear to possess a unique combination of somies. This plasticity was related to hybridization and was not driven by the irradiation per se, since 4 independent sub-clones and one uncloned line of the irradiated L747 parent did not show any changes in somy from the untreated line (Fig. 2B).

For all intra- and interspecific hybrids analyzed, we could observe bi-allelic inheritance of almost all of the homozygous SNPs that are different between the two parental lines (Fig. 2D), indicating that they are full genome hybrids. The SNP inheritance profiles are depicted as circos plots showing the relative parental contributions of homozygous SNPs across all 35 (chr 20 in *L. braziliensis* is a single chromosome product of a fusion between chr 20 and chr 34 but is analyzed as chrs 20.1 and 20.2 in the plot) or 36 chromosomes. While the tetraploid hybrids show generally balanced contributions of chromosomes from both parents, in many cases asymmetric contributions are observed, e.g. chrs 12, 16, 21, 30 in IMA (*Lt*MA37 x *Li*LLM320) hybrid Ac5a, for which three copies are contributed by the *Lt*MA37 parent, and one copy by *Li*LLM320. Rare instances of uniparental inheritance involving a particular chromosome are also observed, e.g. chr 12 in Mondo06 (*Ld*Mongi x *Ld*SL2706) hybrid Cc5a, for which loss of heterozygosity is inferred. There are a substantial number of chromosomes for which the asymmetric or uniparental inheritance patterns involve only part of the chromosome, e.g. chr 32 in IMA Ac5a; chr 17 and 26 in IMA Ba4a; chr 4, 13, 34 in Mondo06 Cb6a; chr 20, 29 in MIR (*Lb*RicX x *Lb*M1) 1a; and chr 14, 32 in MIR 3a, inferring recombination between homologous chromosomes.

### Single cell RNA-sequencing reveals a unique transcriptomic landscape in parasites exposed to γ-radiation

Although expression of protein-coding genes in kinetoplastids is not regulated at the level of transcription initiation, transcriptomic analyses of kinetoplast populations, including *Leishmania*, have revealed clear differences in transcript levels comparing parasites from different life cycle stages or from parasites submitted to stress conditions. These levels are controlled by gene dosage and by a post-transcriptional regulatory network involving different RNA-binding proteins (25, 26). We used single-cell RNA-sequencing (scRNA-seq) between irradiated and non-irradiated promastigotes to help define transcripts and possible rare cell types associated with the sexual mating in *Leishmania*. Using the Chromium Single Cell 3’ workflow (10X Genomics) and Illumina sequencing followed by analysis with the Seurat R toolkit (27), gene expression analysis was carried out on 20,122 and 16,708 cells obtained from duplicate cultures of L747 and MA37, respectively, exposed or not to γ-radiation, one day post inoculation of a log-growth culture at 5×10^6^ promastigotes per mL. After filtering the data to remove potential cell multiplets, dying cells or cells with poor quality transcripts (see Material and Methods), 20,087 L747 (13,392 untreated plus 6,695 irradiated) and 16,645 MA37 (8,156 untreated and 8489 irradiated) cells remained, with expression of 8,156 different genes detected and a median number of 1,079 genes expressed per cell. Since the approach used corresponds to a 3’Tag based sequencing and there are no data available on *L. tropica* transcripts 3’UTR length, we chose *L. major* as the reference transcriptome, as it has the highest-quality *Leishmania* gene annotations (28) and the genomes of both species are highly similar and syntenic.

To estimate the degree of similarity of the global mRNA expression profiles between the different samples and to evaluate the reproducibility of the replicate samples, we performed a principal component analysis (PCA) using a “pseudo-bulk” quality control method (DESeq2) (29). The PCA plot shows the top two principal components which explain most of the variance between the samples, 79% and 16% for PCA1 and PCA2, respectively, and high reproducibility between the replicate samples. The analysis suggests that L747 and MA37 cells present distinct mRNA expression profiles, and that the L747 transcriptome is altered more by the irradiation than MA37 (Fig. 3A). Cells from the replicate cultures were integrated and UMAP (Uniform Manifold Approximation and Projection) was used to visualize the relationship between individual transcriptomes in two dimensionality, where variation between transcriptomes dictates the distance between cells. For the clustering analysis of untreated and treated L747 and MA37, the 3000 most variable genes were selected (Sup. Fig. 6A) and used for Seurat R package unsupervised clustering with reduction dimensions 1-7 and a resolution of 0.3, which identified seven groups of cells, each containing transcriptionally similar cells (Fig. 3B,C). Analysis of the untreated parasite samples separately from irradiated cells revealed a transcriptomic heterogeneity among promastigotes within the same mid-logarithmic-phase culture, which would not be detectable using bulk-RNA sequencing without prior cell sorting. Clustering of untreated cells identified five and six different cell populations in L747 and MA37, respectively (Supplementary Figure 6B), using the same parameters mentioned above. Transcript markers for procyclic and metacyclic promastigote stages, stress granules and protein synthesis are among the cluster-specific genes found in untreated samples (Sup. Fig. 6C-F). The differential expression of some of these markers is evident between untreated L747 and MA37 (Sup. Fig. 6E), highlighting another level of heterogeneity traceable by scRNA-seq.

**Figure 3.**
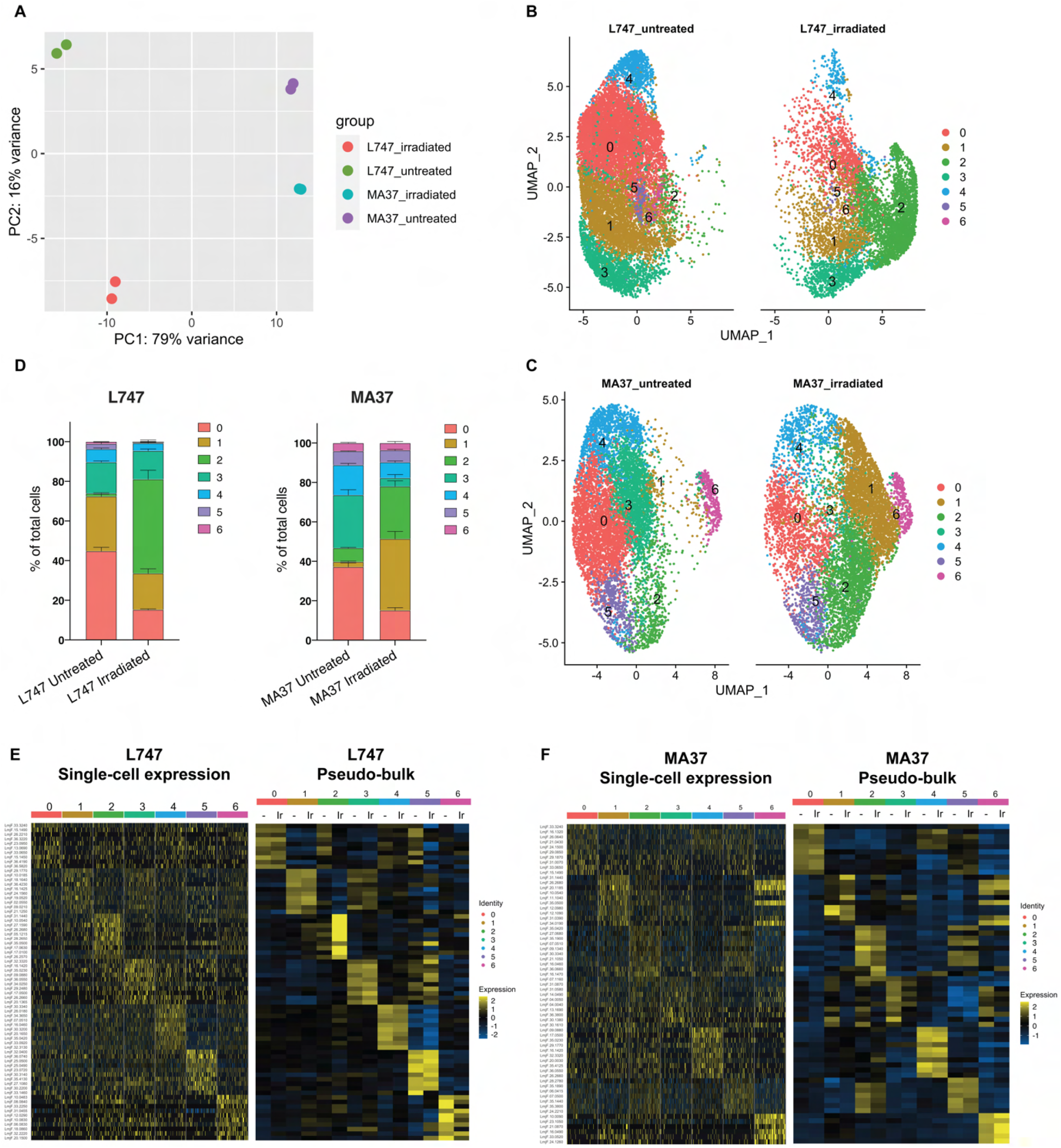
Single cell RNA-sequencing analysis of *L. tropica* parental strains identify discrete clusters of cells expanded after irradiation. (**A**) Principal Component Analysis (PCA) plot of average gene expression comparing scRNA-seq biological replicates of L747 and MA37 promastigote cultures one day post exposure to 6.5 Gy of γ-radiation or not treated (n = 2 for each group). (**B-C**) Unifold Manifold Approximation and Projection (UMAP) visualization of the seven heterogeneous clusters of cells identified in L747 (B) and MA37 (C) according to their transcriptomic profiles using the Seurat R package(27). Cluster identities in L747 and MA37 are completely independent and unrelated with each other. Data presented are a combination of two biological scRNA-seq replicates for each group. (**D**) Representation (%) of each cell cluster per sample in irradiated or untreated L747 and MA37 in the two replicates. (**E-F**) Left panel: Heatmap representation of the expression (log2 of fold change comparing a single cell versus all other cells within the same sample) of the top 10 transcript markers in each cluster identified for L747 (E) and MA37 (F), downsampled to 100 cells in each cluster for visualization (Wilcoxon Rank Sum test adj. *p*<0.05). Right panel: A pseudo-bulk analysis shows the average gene expression of the same top 10 markers, highlighting the differences and similarities between untreated (-) and irradiated (Ir) samples using data from all cells assigned to each cluster.

Next, we analyzed irradiated and untreated cells together, separating the two strains. For both L747 and MA37, submitting the cells to γ-radiation altered the relative distribution of cells within particular clusters, with greatly increased cell numbers in clusters 2 and 1 for the irradiated L747 and MA37 cells, respectively (Fig. 3D). By contrast, the relative abundance of cells in MA37 cluster 3 decreased after treatment, suggesting cells from this cluster might be closely related to the irradiation-induced MA37 cluster 1. The dominant cluster 2 in the irradiated L747 culture stands out as a spatially separated cluster, consistent with the PCA plot showing the relatively strong effect that the γ-radiation has on altering the transcriptome of this strain.

Relative expression of the top 10 upregulated transcripts in each of the seven clusters in each strain revealed the different gene expression profiles in the subgroups of cultured promastigotes (Fig. 3E,F and Sup. Table 2). Despite the fact that the UMAP visualizations failed for the most part to resolve the populations into well separated clusters, the heatmaps showing the different gene expression levels still reveal significant transcriptomic heterogeneity between each of the groups (Fig. 3E,F). A pseudo-bulk analysis of the scRNA-seq data shows differential expression of the top 10 markers between irradiated and untreated cells only within L747 cluster 2, and within MA37 clusters 1 and 3 (Fig. 3E,F). Other clusters remain largely unchanged upon irradiation.

The observation that cluster 2 in L747 (L-cluster2) and cluster 1 in MA37 (M-cluster1) are largely expanded upon irradiation prompted us to investigate a possible transcriptomic signature shared between these two promastigote populations compared with other cells in the same culture. Gene Ontology (GO) analysis revealed that molecular functions such as mRNA binding and kinase activity are significantly enriched in both L-cluster2 and M-cluster1 among the lists of differentially expressed genes (DEGs; |log2FC|>0.1, FDR<0.05) (Fig. 4A). From the 708 genes upregulated post irradiation in L-cluster2 and 296 genes in M-cluster1, 169 are common between the two strains (Fig. 4B, Sup. Table 3). This list includes repressor of differentiation kinase 2 (*RDK2*; LmjF.31.2960) and RNA-binding protein 5 (*RBP5*; LmjF.09.0060), both of which are involved in *Trypanosoma brucei* proliferation (30, 31). Among the top 20 most upregulated genes in either L-cluster2 or M-cluster1 are seven genes common to both lists: serine/threonine-protein kinase *NEK15* (LmjF.26.2570), adenylate-cyclase *ACP2* (LmjF.10.0540), three putative surface antigen proteins (LmjF.05.1215; LmjF.12.1090 and LmjF.35.0550), one gene of unknown function (LmjF.26.2680) and a putative leucine-rich repeat (LRR) protein annotated as a pseudogene on TriTrypDB (LmjF.31.1440) (Fig. 4B). Among the top 20 most downregulated genes in either cluster that are also downregulated in the other cluster are four genes: two ribosomal proteins (LmjF.04.0950 and LmjF.26.1640), a P-type ATPase (LmjF.18.1510) and translation elongation factor 2 (LmjF.36.0190).

**Figure 4.**
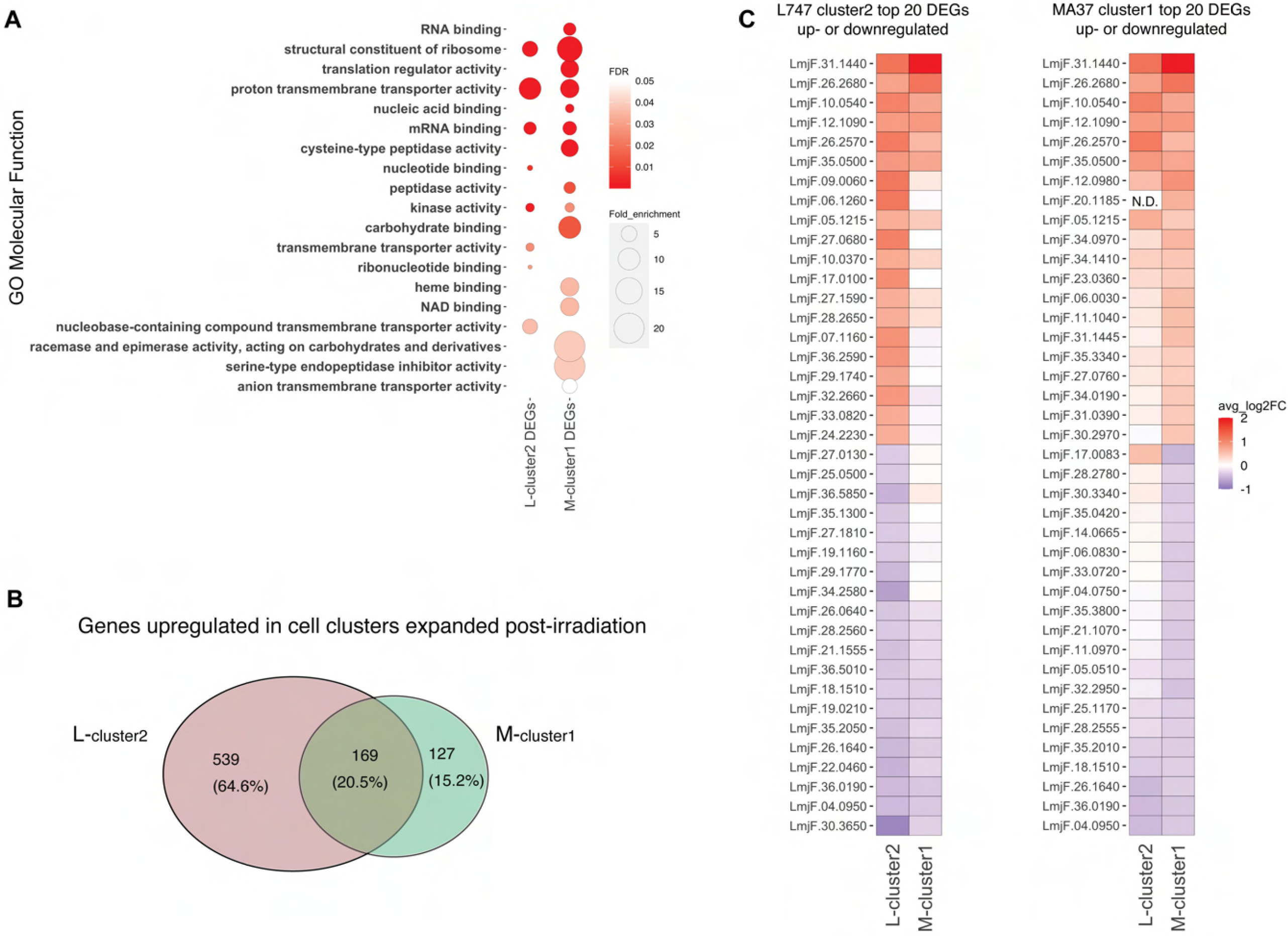
Irradiation-induced cell clusters in L747 and MA37 share a common transcriptomic signature. (**A**) Gene Ontology (GO) Molecular Function enrichment analysis was performed for the lists of differentially expressed genes (DEGs; |log2FC|>0.1; Wilcoxon Rank Sum test adj. *p*<0.05) in L747 cell cluster 2 (L-cluster2) and MA37 cell cluster 1 (M-cluster1) using TritrypDB online tools (Benjamini-Hochberg FDR<0.05). (**B**) Venn diagram representation of the total number of genes upregulated in L-cluster2 and/or M-cluster1 compared to the other clusters (log2FC>0.1; Wilcoxon Rank Sum test adj. *p*<0.05). (**C**) Heatmap representation of the expression of the top 20 genes up- or down-regulated in L-cluster2 (left panel) or M-cluster1 (right panel). Data are presented as the average gene expression in cells from each of the clusters compared with the other clusters in the same sample. Genes are ranked according to their mean expression in both clusters. Expression of gene LmjF.20.1185 was not detected (N.D.) in the L-cluster2 cells.

### Homologues of meiotic genes are enriched in unique clusters of irradiated L747 and MA37 cells

We looked at the expression of genes expressed by the irradiated cells that are potentially involved in genetic exchange. From the list of 169 upregulated genes shared between L-cluster2 and M-cluster1, we identified three putative meiotic gene homologues: a gene encoding the ancestral gamete fusogen HAP2/GCS1 (hapless 2/generative cell specific 1; LmjF.35.0460), the nuclear membrane protein GEX1 (gamete expressed 1; LmjF.04.0100) required for nuclear fusion during yeast mating, and RAD51 recombinase (LmjF.28.0550), involved in DNA-damage repair and genetic exchange in *Trypanosoma cruzi* (32). The average expression of these and additional meiotic gene homologues was also tested by comparing irradiated and untreated cells within each parental line. HAP2, GEX1 and RAD51 are specifically upregulated in the irradiated L747 and MA37 cells, being expressed in 28.4% and 12% of the treated cells, respectively, compared11.9% and 5.75% of the untreated (Fig 5A,B). The single-cell relative expression of HAP2 and GEX1, depicted by UMAP and violin plots, is significantly enriched in L-cluster2 and M-cluster1 in irradiated samples (Fig. 5C-F) compared to the other cell clusters in any condition.

**Figure 5.**
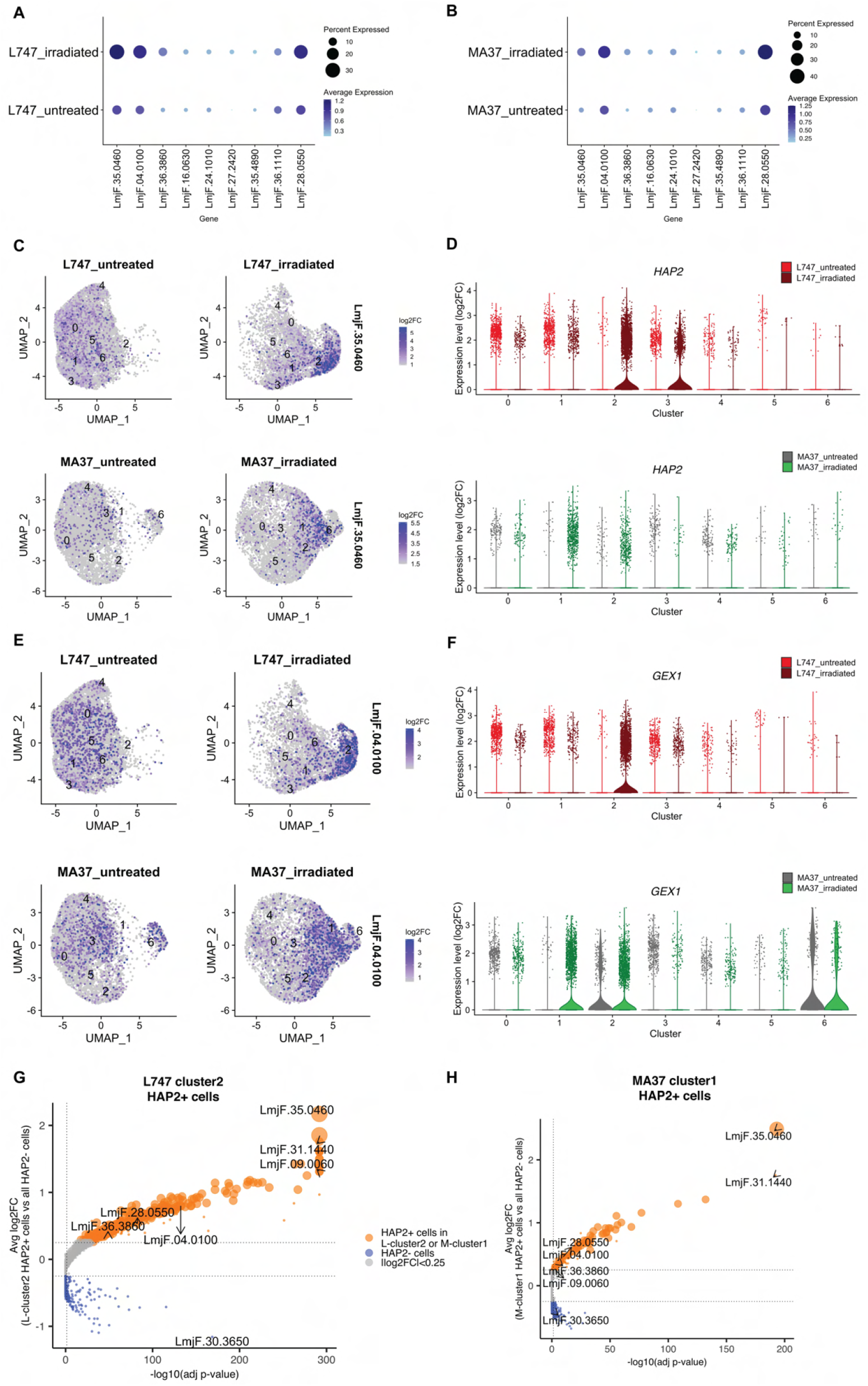
Expression of HAP2 and GEX1 is enriched in irradiation-induced L747 cluster 2 and MA37 cluster 1. (**A-B**) Dotplots depicting the average expression of *Leishmania* homologues of genes involved in genetic exchange and meiosis in other organisms, shown for L747 (A) and MA37 (B) populations with and without exposure to irradiation (pseudo-bulk analysis). The dot size in the plots represents the percentage of cells expressing those genes in each sample. Meiotic genes tested are, from left to right: *HAP2* (LmjF.35.0460), *GEX1* (LmjF.04.0100), *HAP2-like* (LmjF.36.3860), *SPO11* (LmjF.16.0630), *MND1* (LmjF.24.1010), *HOP2* (LmjF.27.2420), *DMC1* (LmjF.35.4890), *HOP1* (LmjF.36.1110) and *RAD51* (LmjF.28.0550). (**C-E**) UMAP plots showing the expression of HAP2 (C) and GEX1 (E) in the different cell clusters identified in L747 (upper panels) and MA37 (lower panels) untreated or irradiated. (**D-F**) Violin plots representation of the single-cell expression data of HAP2 (D) and Gex1 (F) in L747 (upper panel) and MA37 (lower panel), comparing cells from untreated and irradiated cultures (**G-H**) Volcano plot representation of DEGs in HAP2^+^ cells (i.e., cells with HAP2 expression log2FC>1.0) identified in L-cluster2 (G) and M-cluster1 (H). Delta percent expressed (%HAP2^+^ minus %HAP2^-^ cells expressing a particular gene) is presented as different data point sizes in the plots. Threshold values for adjusted *p*<0.05 (vertical dotted line; Wilcoxon Rank Sum test) and |log2FC|>0.25 (horizontal dotted lines) were used.

### HAP2 expression is associated with *L. tropica* mating competence *in vitro*

The well conserved class II fusion protein HAP2 is involved in the fusion of gametes or opposite mating type cells in many organisms (33). Expression of HAP2 has been previously used to identify gametes and immediate precursors in the related trypanosomatid *Trypanosoma brucei* found inside the salivary glands of the tsetse fly vector (34). We hypothesized that *L. tropica* HAP2 might be a marker for mating competent cells. First, we investigated the list of genes co-expressed with *HAP2* in the irradiation-induced M-cluster1 and L-cluster2 (*HAP2*^*+*^ cells vs the total *HAP2*^-^ cell population; |log2FC|>0.25, adj. *p*<0.05). Several genes that were upregulated in cells from L-cluster1 and M-cluster2 as a whole (Fig 4c), e.g. *RBP5* (LmjF.09.0060) and LmjF.31.1440, were also upregulated in the *HAP2*^+^ cells within these clusters, as were other meiosis-related genes that are common to these clusters; *GEX1* (LmjF.04.0100), *RAD51* (LmjF.28.0550), and LmjF.36.3860, encoding a HAP2-like protein.

In order to follow the expression of HAP2 at the protein level, we generated reporters of expression by tagging the endogenous protein with a fluorescent mNeonGreen (mNG) fusion in both L747 (Fig. 6A) and MA37 (Fig. 6F). In each case the tag was inserted by CRISPR/Cas9 in the N-terminal part of the ORF, so that the endogenous sequences, in particular the 3’UTR involved in post-transcriptional regulation, would not be affected. The intensity of mNG expression detected by flow cytometry was low in both parental strains, but in both cases a proportion of cells reproducibly expressed a fluorescence level higher than the negative control (L747 or MA37 cultures without the reporter construct; Fig. 6B,G). In L747 cultures, an average of only 3.0% of cells expressed mNG-HAP2 in the untreated condition at day 1 post inoculation, whereas at the same time point 23.4% of the L747 irradiated cells expressed mNG-HAP2 (Fig. 6C). By day 2 post-inoculation, mNG-HAP2^+^ cells in the irradiated cultures were detected at frequencies closer to those of untreated cultures (Fig 6C). A similar pattern of increased frequency of mNG-HAP2^+^ cells was observed for L747 parasites treated with either H_2_O_2_ or MMS, although in each case the elevated expression persisted until day 4 of culture, and the effect was especially pronounced in the MMS-treated cells (60.2% mNG-HAP2^+^). For the MA37 reporter line, the proportions of mNG-HAP2^+^ cells were not significantly different at day 1 of culture (32.8% of the untreated and 28.9% of the irradiated parasites, *p =* 0.8857 Mann-Whitney test), which was consistent in all stress treatments (Fig. 6H-J). By day 2, the relative frequencies were altered, with levels declining in the untreated cells and increasing in the irradiated cells (Fig. 6H). Similar patterns of mNG-HAP2 expression were observed in cells treated with either H_2_O_2_ or MMS (Fig. 6I,J), with MMS inducing a more long-lasting effect. The relatively high frequency of mNG-HAP2^+^ cells in the untreated MA37 cultures was surprising given the scRNAseq comparisons showing the large expansion of HAP2 expressing cells in the irradiation-induced cluster, and suggests that HAP2 can be expressed under non-stress conditions even if not necessarily within the same population of promastigotes. By contrast, HAP2 expression in the L747 parent appears more dependent on induction by exposure to stress. The HAP2 expression profiles of the reporter constructs seem to more accurately reflect the pseudo-bulk RNA-seq comparisons (Fig 5A), which show a relatively small difference between irradiated and untreated MA37 cells, and a large difference between irradiated and untreated L747 cells.

**Figure 6.**
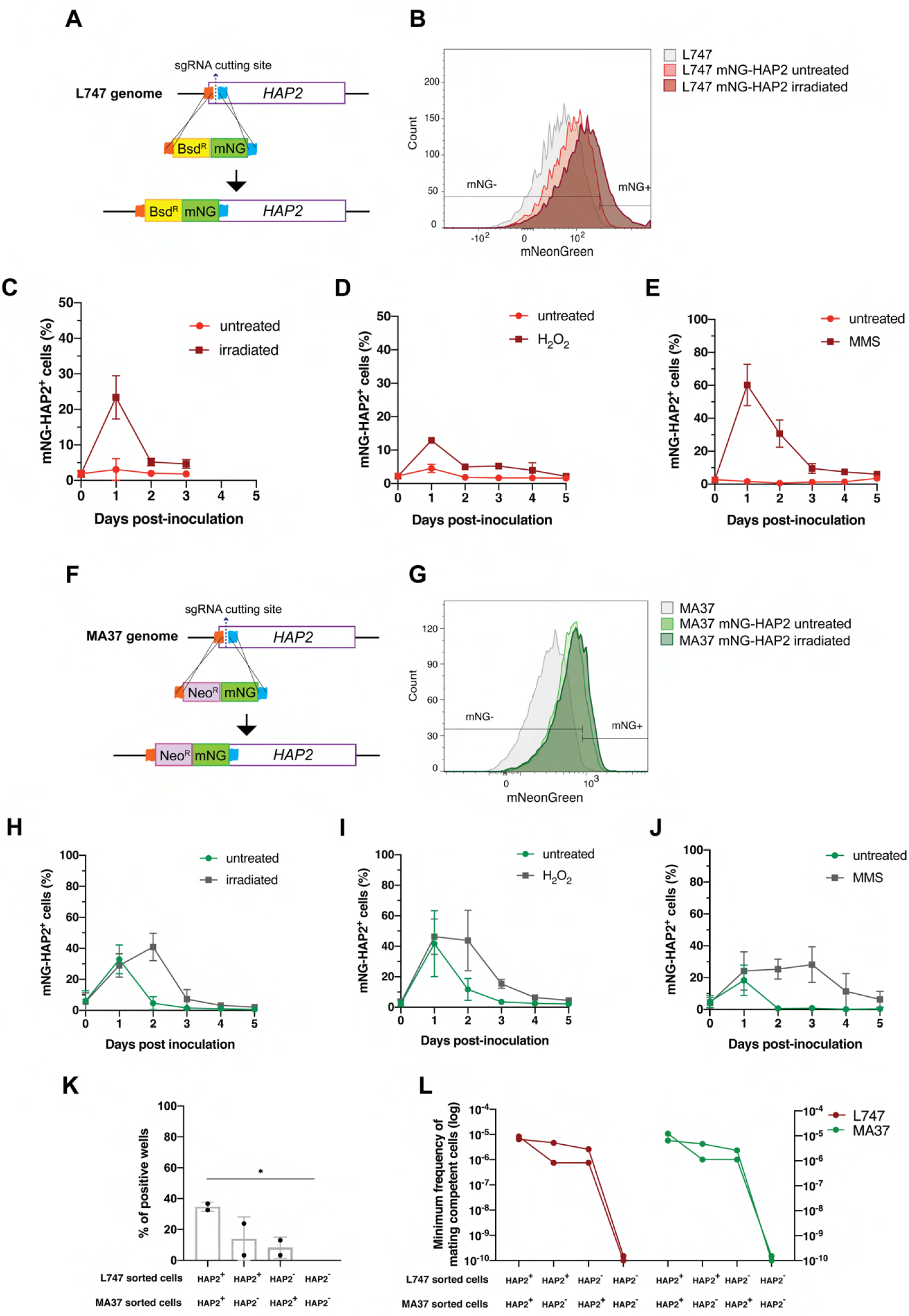
HAP2 protein expression in *L. tropica* promastigotes is associated with *in vitro* mating capacity. (**A-F**) Schematic representation of the mNeonGreen (mNG)-HAP2 fusion reporter construct generated in the L747 strain (A) and in the MA37 strain (F). (**B-G**) Flow cytometry histogram plots showing the fluorescence intensity of mNG-HAP2 in L747 (B) and in MA37 (G) one day post-inoculation, with or without irradiation. (**C-E**) Proportion of mNG-HAP2^+^ cells over time in L747 mNG-HAP2 cultures with or without exposure to γ-radiation (C), to H_2_O_2_(D) or to MMS (E). (**H-J**) Proportion of mNG-HAP2^+^ cells in MA37 transfectants after exposure to γ-radiation (H), to H_2_O_2_ (I) or to MMS (J). (**L**) Proportion of culture wells exhibiting growth of LMA hybrids after *in vitro* crosses using the different indicated pair-wise combinations of FACS-sorted mNG-HAP2^+^ and mNG-HAP2^-^ cells one day after irradiation. \**p* = 0.0440 (one-way ANOVA, Tukey’s test). (**M**) Minimum frequency of mating competent cells for both L747 and MA37 parental strains in *in vitro* crosses using different combinations of FACS-sorted mNG-HAP2^+^ and mNG-HAP2^-^ cells (n = 2 independent experiments). Frequencies collected from each single experiment are linked by a line. Red lines and green lines represent data from L747 and MA37 parental strains, respectively.

We submitted the parental mNG-HAP2 reporter lines to γ-radiation and purified mNG-HAP2^+^ and mNG-HAP2^-^ cells by fluorescence-activated cell sorting (FACS) one day post-irradiation (Fig. 6B,G), and co-cultured different combinations of the selected populations. Due to technical restrictions in the number of HAP2^+^ cells that could be sorted, approximately 10 fold fewer promastigotes of each parent were used in these crosses compared to those performed using unsorted cells (Sup. Table 1). Co-cultures that were established using the mNG-HAP2^+^ cells from both parents produced the highest frequency of hybrids (34% positive wells), with a minimum frequency of mating competent cells 7.5 fold the frequency detected in crosses using unsorted cells (Figs. 6K,L and 1D; Sup. Table 1). Interestingly, the presence of just one parental HAP2^+^ cell population was sufficient to generate hybrids, albeit at lower frequencies (16% and 11% positive wells; 0.2-0.37 fold the minimum frequency of mating competent cells versus the double positive pair). Critically, co-cultures seeded with the mNG-HAP2^-^ cells from both parents failed to generate any hybrids.

## Discussion

Reproduction in *Leishmania* includes a cryptic sexual cycle that operates among the extracellular promastigote stages developing in the sand fly vector. The mating competent cells, and in particular the existence of a gametic stage, have yet to be identified. Studying the biology of mating in *Leishmania* would be greatly facilitated if these events could be reproduced using cells from axenic cultures, and we have recently achieved the recovery of stable hybrids generated between cultured promastigotes entirely *in vitro* (16). While these initial findings established that mating competent forms can arise in axenic culture, the mating frequencies were extremely low, and only a single pairwise combination of *L. tropica* parental lines would successfully hybridize. In the current studies, we demonstrate that under various conditions of environmental stress, including exposure to H_2_O_2_, methyl methanesulfonate, or γ-radiation, the *in vitro* mating capacity of the *L. tropica* strains could be markedly enhanced and extended to other *L. tropica* strains and multiple *Leishmania* species, including *L. infantum, L. donovani*, and *L. braziliensis*. The products of these mating events were in each case full genome, polyploid hybrids. By single cell RNAseq analysis, we could identify transcriptionally unique populations of promastigotes whose numbers were drastically increased in each parental line following exposure to γ-radiation, and for which specific meiotic gene homologues, including the ancestral gamete fusogen HAP2, were found to be upregulated. By generating reporter constructs for HAP2, we could select for promastigotes that could hybridize or not *in vitro*.

Our findings add to the list of facultatively sexual eukaryotes, including yeast and green alga, that exhibit more sexual events under stress conditions. As each of the stress conditions that we imposed are known to produce DNA damage, the results also support studies in varied model eukaryotes suggesting that DNA repair is a central adaptive function of meiotic recombination (17-19). Homologous chromosome pairing during meiosis provides an undamaged, template chromosome for DNA repair. The generation and repair of double-strand breaks is evidenced in our *in vitro* hybrids by the substantial number of chromosomes for which there were changes within the chromosome in the relative contribution of the parental alleles. A role for SPO11 protein in the induction of programmed DNA double-strand breaks is argued to be a universal feature of meiotic crossover events in eukaryotes (35). However, meiotic recombination induced by X-irradiation in *S. cerevisiae* and in *C. elegans* was shown to occur even when SPO11 was absent or non-functional (23, 24). In our studies, expression of the homologous *SPO11* gene in *L. tropica* was not upregulated in the irradiated, mating competent cells.

In order to identify mating competent cells present in *Leishmania* cultures, we performed single cell RNA sequencing of L747 and MA37 cultured cells exposed or not to γ-radiation. To our knowledge, this is the first application of single cell transcriptomics in *Leishmania*, although it has been applied to other kinetoplastid pathogens to reveal the heterogeneity of the parasite populations arising in their mammalian hosts or insect vectors (reviewed in (36). Our analyses revealed transcriptomic heterogeneity within and between the parental *L. tropica* lines in the promastigote cultures. The heterogeneity likely reflects cells in different phases of the cell cycle, as well as differences in the developmental stage progression of the cultured promastigotes. The extensive differences in the transcriptomes of cells comparing the parental lines, apparent in both the PCA and UMAP visualizations, is surprising given the genetic similarity between these *L. tropica* strains (26), and likely reflects their different culture histories. More meaningful to the current studies is the cluster heterogeneity comparing the untreated versus irradiated cells, which for each parental line revealed the emergence of a transcriptionally unique population in the irradiated cultures. Probing the clusters for enriched transcripts related to meiotic processes identified homologues of HAP2, GEX1, and RAD51, involved respectively in cell fusion, nuclear fusion and DNA repair, that were largely confined to the unique clusters that emerged in the irradiated cultures. Even though a large fraction of the transcriptional shift is likely explained as part of a more global cellular mechanism to recover from stress caused by the irradiation, the expansion of cells expressing HAP2 transcripts in treated samples, the co-regulation of other meiotic genes (e.g., a HAP2-like FusM homologue and GEX1) and validation *in vitro* using HAP2 reporters, strongly suggest that these cells are associated with the enhanced mating frequency observed post-treatment. The possible role in genetic exchange of other genes coordinately expressed with HAP2 but without any described meiotic function will be investigated in future studies. Of note, RNA-binding proteins, such as RBP5 and ZC3H11, were found to be upregulated in these cells and could have novel regulatory functions during *Leishmania* mating.

HAP2 is a type I transmembrane protein that displays the same three-dimensional fold as class II viral fusion proteins. It plays a key role in the membrane fusion of gametes or mating cells that was first identified in plants, and is broadly conserved from protists to invertebrates (33, 37-40). HAP2 expression was recently reported in *T*.*brucei*, both in gametes and in meiotic intermediates (34). In the current work, we generated reporter constructs for HAP2 in L747 and MA37 and could show enhanced expression or altered expression kinetics in the stressed cultures. By sorting the cells into mNG^+^ and mNG^-^ populations, we could confirm that HAP2 expression is a marker for mating competent cells since hybridization failed unless the co-cultures included mNG^+^ selected cells from one or both of the parents. To the extent that HAP2 is itself required for hybridization, then the apparent requirement for HAP2 expression on just one of the fusing cells is consistent with a number of studies showing that it can function unilaterally; either on the mating type minus gamete in *Chlamydomonas* algae (33, 39), or on the male gamete in *Plasmodium* (39), the ciliated protozoan *Tetrahymena thermophila* (37), or the plant *Arabidopsis thaliana* (38, 40). It is interesting that the MA37 parent showed a relatively high frequency of HAP2 expressing cells even absent exposure to the exogenous stress conditions. This may explain why this *L. tropica* strain hybridizes with such high efficiency in flies (12), and why we are able to obtain some *in vitro* hybrids even without exogenous stress so long as MA37 is a mating partner.

Contrary to the *in vitro* LMA hybrids that were previously generated, which were diploid, triploid and tetraploid (16), the irradiation-facilitated LMA hybrids, as well as the other intra- and interspecies hybrids described in this report, are in their great majority tetraploid. These results strongly suggest fusion between the two diploid parental cells, and that the stress conditions that promoted the fusogenic potential of the cells may have produced unusual features leading to unreduced gamete formation, which is a common mechanism underlying polyploidization in plants (41). Attempts to condition the tetraploid hybrids to complete a putative meiotic program by passing them through flies or by re-exposure to stress conditions failed to reduce their ploidy (Sup Figure 5), showing that their polyploid state is relatively stable. As tetraploid progeny are occasional products of hybridization events in flies, it is also possible that tetraploid intermediates, as described for some fungi (42, 43), are a normal part of the mating process in *Leishmania*, with the ploidy reduction steps somehow lacking in the *in vitro* protocol. Since in the related protist *T. brucei*, haploid gametic cells have been identified prior to fusion in cells localized to the salivary gland of the tsetse fly vector (44), we favor a comparable sexual process can operate during *Leishmania* development in sand flies. The fact that a few triploid hybrids, also observed in flies, were generated by the *in vitro* protocol, could mean that a haploid gamete was produced by at least one of the parents, and may encourage additional manipulations of the culture conditions to promote a more complete and uniform sexual cycle *in vitro*.

It is interesting that while the hybrids appear unable to undergo the ploidy reductions associated with meiotic divisions, they seem to have retained an ability for homologue pairing and recombination after fusion, as evidenced by the breaks within chromosomes in the relative contributions of the parental alleles. Since the maintenance of the tetraploid state (two genome complements from each parent) would be expected to mask recombinations generated by reciprocal crossover events typically associated with meiosis I, the data suggest noncrossovers or gene conversion as a mechanism of DNA repair that would result in gene exchange in only one of the homologs. The asymmetric parental contributions observed both within and between different chromosomes could also have arisen by chromosome loss or gain at the level of individual somy, which is well described for *Leishmania* genomes during vegetative growth (45, 46). Aneuploidy, including mosaic aneuploidy, is a constitutive feature of the *Leishmania* genome (47). Since gene dosage can directly control the levels of gene expression in *Leishmania* and impact phenotypes, (26, 48), the chromosome-specific patterns of aneuploidy and recombination in the hybrids can be used for linkage studies. It is notable that the progeny that were recovered are in almost every case distinct from the other progeny clones generated in the same experimental cross with respect to the combinations of the relative parental contributions to individual chromosomes that they display. The distinct haplotypes observed in each of the 17 hybrids generated between the *L. donovani* strains from India and Sri Lanka are a case in point. As these parental strains produce, respectively, visceral and cutaneous forms of disease in humans and in mice (49, 50), it may be possible to map the genes controlling the behavior of the progeny clones in mice by genome wide association studies.

The stress-facilitated, *in vitro* hybridization protocol described in this report removes the requirement for sand flies as the main constraint to generating large numbers of recombinants between *Leishmania* species and strains, and offers a transformative approach for genetic analysis to understand the extraordinary phenotypic diversity of the genus.

## Material and Methods

### *Leishmania* cultures and transfection

Promastigotes were grown in axenic cultures at 26°C in complete medium 199 (CM199) (13). The following parental *Leishmania* lines have been previously described (12): *L. tropica* L747 RFP-Hyg (MHOM/IL/02/LRC-L747), *L*.*tropica* MA37 GFP-Neo (MHOM/JO/94/MA37), *L*.*major* LV39 RFP-Hyg (MRHO/SU/59/P-strain), *L*.*major* FV1 Sat (MHOM/IL/80/Friedlin) *L*.*infantum* LLM320 RFP-Hyg (MHOM/ES/92/LLM-320; isoenzyme typed MON-1). The following additional parental lines were generated for the purposes of this study: *L*.*tropica* Moro RFP-Hyg (MHOM/AF/19/Moro), isolated from a cutaneous lesion in a patient from Afghanistan; *L*.*donovani* Mongi RFP-Hyg (MHOM/IN/83/Mongi-142), isolated from a patient with visceral leishmaniasis in India; *L*.*donovani* SL2706 GFP-Neo (MHOM/LK/19/2706), isolated from a cutaneous lesion in a patient from Sri Lanka; *L*.*braziliensis* M1 GFP-Neo or RFP-Hyg (MHOM/BR/00/BA779), isolated from a cutaneous lesion in a patient from Brazil, and *L*.*braziliensis* RicX GFP-Neo or RFP-Hyg (MHOM/PE/19/RicX), isolated from a cutaneous lesion in a patient from Peru.

For the parental lines generated in this study, *Swa*I-digested pA2-GFP-Neo or pA2-RFP-HYG plasmids (51) were gel purified and the linear fragment flanked by the small subunit rRNA (SSU) homology regions was transfected into log-phase promastigotes using an AMAXA Nucleofector 4D (Lonza). Parasites (8 × 10^6^ cells) were harvested, washed in PBS and resuspended in 20 µL supplemented P3 primary cell buffer (Lonza). DNA fragments (0.5-1 µg) and cells were mixed in a Nucleofector Strip and transfection was performed immediately using program FI-115. Parasites were incubated at room temperature for 2 min in 100 µL CM199 and for 16 h in 5 mL CM199 before selection of positive transfectants using appropriate drug pressure (see “*In vitro* generation of hybrids” below).

### *In vitro* generation of hybrids

*Leishmania* hybrids were generated following the protocol previously developed (16). Briefly, cultures of parental strains carrying resistance to a different antibiotic marker were initiated in parallel. One day post inoculation, the concentration of each culture was estimated by counting under a hemocytometer and equal volumes of the parental cultures were mixed together and immediately distributed in 96 well plates (100 µL/ well). The number of cells distributed per well varied according to the experiment, and are summarized in Sup. Table 1. The cocultures were transferred into a selective medium containing both antibiotics in 24 well plates on the next day (100 µL in 1mL). The following antibiotics were used for double-drug resistant hybrid selection: Geneticin (Neomycin analog; Thermo Fisher; Neo 50 µg/mL), Hygromycin B (Sigma-Aldrich; Hyg 25 µg/mL), Nourseothricin (Fisher scientific; Sat 200 µg/mL) and Blasticidin S (Fisher Scientific; Bsd 20 µg/mL). The hybrid nature of the double drug resistant lines was by verified by their co-expression of fluorescence markers, analyzed by flow cytometry using a FACSCanto II system and FACSDiva software (BD Biosciences) (Sup. Fig. 2). For hybrids generated from parents that did not carry different fluorescence markers, the presence of both parental antibiotic resistance markers was validated by PCR on DNA extracted from hybrid clones (DNeasy blood and Tissue Kit, Qiagen), targeting the antibiotic resistance genes (see Sup. Table 4 for the sequence of PCR primers). For testing the effects of stress inducing conditions on the generation of hybrids, parental cultures were initiated in parallel to untreated cultures and immediately exposed to 6.5 Gy of γ-radiation or supplemented with 250µM of Hydrogen Peroxide (H_2_O_2_) or 0.005% of Methyl Methane Sulfate (MMS). 4 repetitions of each experiment were performed. For graphic representation on a logarithmic scale of the minimal frequencies of mating competent cells in each parental culture (Figures 1 and 5), a minimal value of 1e-10 was added to each frequency so the experiments where no hybrids were obtained.

### DNA staining, flow cytometry and confocal microscopy

To estimate the ploidy of *Leishmania* hybrids, promastigotes at logarithmic growth phase were stained in propidium iodide (PI) followed by flow cytometry analysis, as previously described (16). Briefly, cells were fixed in 0.4% paraformaldehyde for 1 min at room temperature, sedimented, resuspended in 100 µL PBS and permeabilized in 1mL methanol for 15 min on ice. After centrifugation, cells were incubated in 1 mL PBS for 10 min at room temperature. Next, DNA was stained using 500 µL of a PI-RNase mix (13 mg/mL each) at room temperature for 10 min. Stained cells were washed in cold PBS and analyzed on a FACSCanto II (BD Biosciences) using FACSDIVA software. Flow cytometry data were analyzed using FlowJo software v.10.7 (Becton Dickinson & Company).

For microscopic visualization of nuclei and kinoteplasts by DNA staining, parasites were fixed in 4% paraformaldehyde, centrifuged and resuspended in 0.5% Glycine and incubated 10min at room temperature. They were then pelleted again, washed in PBS and incubated 30min in 10µg/mLHoechst 33342 (Thermo Fisher). The fixed parasites were then mounted on polylysine-coated glass microscope slides and images were acquired on a SP8 confocal microscope (Leica microsystems).

### Whole genome sequencing analysis

DNA was purified from log-phase promastigote cultures using the DNeasy Blood and Tissue kit (Qiagen) according to manufacturer’s instructions and submitted to Psomagen for next-generation sequencing (Rockville, MD). DNA libraries were generated using TruSeq Nano DNA Library Prep kit (Illumina) and the 100bp-paired-end reads were sequenced on a HiSeq2500 (Illumina). Fastq files were quality checked with FastQC software. Read trimming and filtering were performed with Trimmomatics (52), resulting in > 7.8M reads per library. Mean sequencing coverage in the dataset was 33.5, according to Qualimap v.2.2.1 (Sup. Table 5). Paired-end reads were aligned to the closest reference genomes available from TriTrypDB (tritrypdb.org; *L. tropica* L590 v.50, *L. donovani* BPK282A1 v.50, *L. infantum* JPCM5 v50, *L. braziliensis* M2904 v.50) using BWA v.0.7.17 with default parameters. The PAINT software suite (53)was used to find and extract the homozygous SNP marker differences between each parental strain pair. The alleles and their frequencies in the different hybrid progenies were found using the getParentAllelFrequencies utility in PAINT. Chromosome somies were determined by calculating the normalized read depth with the ConcatenatedPloidyMatrix in PAINT. In the case of polyploid hybrids, the somy values were divided by 2 and multiplied by the estimated ploidy from the DNA content analysis and the parental contribution profile. Somies were reported as heatmaps generated using the gplots and ggplot2 packages in R. Circos plots of the inferred parental contribution were generated using the Circos software (54).

### Single-cell RNA-seq library preparation and sequencing

Promastigotes at early-log phase were treated with 6.5 Gy of γ-radiation and compared with untreated cultures using scRNA-seq. Cells were harvested by centrifugation one day post-irradiation/inoculation and samples were prepared according to Chromium Next GEM Single Cell 3’ Reagent Kits v3.1 (10X Genomics) manufacturer’s instructions. Briefly, 5×10^3^ cells were loaded onto the 10X Chromium Controller microfluidics system and combined into microdroplets with a unique bead carrying oligonucleotides containing an Illumina adapter, a 10x cell barcode, a Unique Molecular Identifier (UMI) and a poly(dT) sequence. Co-partitioned cells were submitted to lysis, followed by reverse transcription of poly-adenylated mRNA, cDNA amplification, enzymatic fragmentation and size selection. Final cDNA library quality control was assessed using a 4200 TapeStation system (Agilent) and Qubit fluorometer and dsDNA HS Assay Kit (Invitrogen). Libraries were submitted to Psomagen (Rockville, MD), pooled and sequenced using two lanes of HiSeqX (Illumina; 150bp paired-end; read length 28*8*91). Two biological replicates of each *L. tropica* culture were sequenced.

### scRNA-seq data pre-processing and quality control

Raw sequencing data were processed using Cell Ranger software v.5.0 (10X Genomics). Sequence reads were demultiplexed into fastq files using Cell Ranger *mkfastq*. Reads were quality checked using FastQC, trimmed and filtered using Fastp for average read quality of at least 20 and minimum length of 20 bp. Since Cell Ranger currently does not provide a built-in reference *Leishmania* transcriptome, a custom reference was generated using *L. major* Friedlin genome v.50 fasta and gff annotation files from TritrypDB. Reliable 3’UTR annotation was necessary for appropriate mapping of the scRNA-seq reads, as the 3’tag based approach used here only targets the 3’end region of transcripts and eventual polyA stretches in the middle of genes. Therefore, absence of 3’UTR annotation available for *L. tropica* strains made it necessary to use the mapped 3’UTR coordinates from genetically close *L. major* (28). Using custom scripts, the *L. major* transcript coordinates were included in the *L. major* gff annotation file were extended to include the predicted 3’ ends of transcripts and renamed to “exon” to be read by Cell Ranger *mkgtf* and *mkref. L. major* mitochondrial DNA sequence (kDNA maxicircle) was added to the reference for eliminating the cell barcodes associated with dying/dead cells later. The resulting custom reference transcriptome was used to align filtered reads and quantify the number of different UMIs for each gene using Cell Ranger *count*. Data from irradiated and untreated samples from the same strain were aggregated together using Cell Ranger *aggr*. The fraction of reads containing a valid barcode assigned to a cell and confidently mapped to the transcriptome varied between 87%-94% in MA37 and 85.7%-86.8% in L747 libraries. The estimated number of cells detected in each replicate was between 3,245-7,485 with 20,401-49,580 mean reads per cell (Sup. Table 6). The generated UMI count matrices were loaded into RStudio v.1.4.1717 and further processed using the Seurat R package v.4.0.3 (27). Low-quality cells were removed according to the following parameters: (1) cells in which maxicircle genes represent > 10% of total expression, (2) cells with expression of fewer than 200 different transcripts or (3) more than 5,000 different transcripts. The first two filters were used to account for dying/dead cells and empty beads and the third for multiplets in the same droplet.

### Cell clustering and differential expression analysis

Filtered data were normalized and variance stabilized by regularized negative binomial regression using the *sctransform* utility (55) for cell clustering, dimensionality reduction and Unifold Manifold Approximation and Projection (UMAP) visualization in Seurat. This package provides a fast approach to help decrease possible batch effects in the clustering. Maxicircle gene expression was regressed out of the sctransform analysis to avoid effects on the data clustering. For differential gene expression analyses reported in violin plots, dot plots and heatmaps, the raw UMI counts were normalized by library read depth, log-transformed, centered and scaled (Z-scored). The two replicates of untreated and irradiated samples from the same strain were integrated and analyzed as a single Seurat object. The 3,000 most variable genes were identified using *FindVariableFeatures* utility and used to perform principal component analysis (PCA). Based on empirical data, the top seven principal components were used to build a shared nearest neighbor graph and modularity-based clustering using the Louvain algorithm with a resolution of 0.34 using *FindNeighbors* and *FindClusters*, which resulted in seven clusters for both *L. tropica* strains. UMAP visualization was calculated using these seven neighboring points. Transcript markers for each cluster were identified by comparing the average expression and the percent expressed of each gene in the cells within a cluster against the rest of the cells in the sample using *FindMarkers*. For the pseudo-bulk quality control analysis at the sample level of the scRNA-seq replicates shown in Figure 3a, the DESeq2 R package was used. Briefly, *DESeqDataSetFromMatrix* was used to create a DESeq object using the raw UMI counts, which were then normalized and log-transformed using *rlog* to account for differences in library depth and transcript composition. PCA plots were generated to visualize the first two principal components that best explain the variance in the data.

### HAP2mNG protein expression reporters

The L747-mNG-HAP2 and MA37-mNG-HAP2 reporter lines were generated by inserting resistance markers for Blasticidin and Neomycin, respectively, followed by an N-terminal mNeonGreen tag by CRISPR/Cas9 gene editing, following the strategy previously described (56). The correct insertion of the tag was validated by two PCRs: the first using a forward primer located in the 5’UTR part of the Hap2 gene and a reverse primer located in the middle of the ORF; the second using the same forward primer and a reverse primer located at the end of the resistance marker inserted with the mNeonGreen (see Sup. Table 4 for the complete list of primers used for the CRISPR/Cas9 strategy). The expression of the tagged protein was followed day by day using flow cytometry. At least 3 repetitions were performed for each treatment (6.5 Gy of irradiation, 150 µM of H_2_O_2_ and 0.005% of MMS), made simultaneously with untreated controls. The estimation of the proportion of cells expressing the mNG-HAP2 was estimated using the FlowJo software v10.7 (Becton Dickinson & Company), using identically treated L747 T7Cas9 and MA37 T7Cas9 cultures as negative controls for normalization.

### *In vitro* mating of mNG-HAP2 selected cells

For *in vitro* mating experiments using HAP2^+^-sorted cells, the mNG-HAP2 reporter lines were transfected with *Pme*I-*Pac*I-digested pSSU-tdTomato-Neo (kindly provided by Dr. Deborah Smith, University of York, UK) or SwaI-digested pLEXSY-cherry-Sat2 (EGE-236, Jena Bioscences) following the same procedure described for GFP-Neo and RFP-Hyg above (see “*Leishmania* cultures and transfection”). Irradiated cultures of L747 mNG-HAP2 and MA37 mNG-HAP2 were sorted one day post-inoculation and irradiation based on the expression of mNG, using L747 T7Cas9 and MA37 T7Cas9 cultures as negative controls of fluorescence expression. After sorting, the mNG-HAP2^+^ and mNG-HAP2^-^ cells from each parent were mixed in various combinations and the co-cultures were incubated overnight at 26°C in 96 well plates (100µL/well). The co-culture wells were transferred individually in double-drug selective medium (Bsd + Neo) the following morning. Verification of positive growing hybrids was performed using a BD LSRFortessa system and FACSDiva software (BD Biosciences) to detect cells positive for both mCherry and tdTomato (Sup. Fig. 7). Data were analyzed using FlowJo software v10.7 (Becton Dickinson & Company).

### Transmission electron microscopy

Promastigotes at logarithmic growth phase (10^7^ parasites) were harvested by centrifugation, washed in cold PBS) and resuspended in 1.5 mL of fixative solution (2.5% glutaraldehyde in 0.1 M sodium cacodylate buffer). Fixed cells were submitted to NIAID Electron Microscopy Unit (Hamilton, MT). All subsequent steps were carried out using a Pelco Biowave microwave at 250 Watts under vacuum. After rinsing in buffer, the cells were post-fixed with 1% reduced osmium tetroxide and treated with 1% tannic acid. The cells were stained with uranyl acid replacement stain, dehydrated with ethanol, infiltrated with Epon-Araldite resin, and polymerized overnight in a 60°C oven. Ultrathin sections were cut using a Leica UC6 ultramicrotome and imaged on Hitachi7800 TEM using an AMT camera.

### Statistical Analyses

All statistical analysis for the whole-genome sequencing data and scRNA-seq were performed using specific R packages in Rstudio v.1.4.1717. Graphpad Prism v.8.0 was used for statistical analysis of other data comparisons.

## Data availability

The raw sequence data containing reads from the 51 WGS samples and 8 scRNA-seq samples sequenced are deposited in the SRA database with Accession numbers PRJNA756557 and PRJNA756571, respectively. The data that support the findings of this study are available from the corresponding author upon reasonable request. Summary statistics on the sequencing data are available in Supplementary Tables 5 and 6.

## Acknowledgements

We thank Margery Smelkinson and Owen Schartz (NIAID Biological Imaging Section) for the technical support with confocal imaging; Calvin Eigsti, Thomas Moyer and Iyadh Douagi (NIAID Flow Cytometry Section) for the technical support with flow cytometry data acquisition and cell sorting; Vinod Nair (Electron Microscopy Unit, RML, NIH) for the support with electron microscopy image acquisition and Nilakshi Samaranayake and Hasna Riyal (Faculty of Medicine, University of Colombo) for coordinating the parasite propagation and passage of Sri Lankan isolates. We also thank Nagib El-Sayed (University of Maryland), Steve Beverley (Washington University) and P’ng Loke (NIAID, LPD) for helpful discussions. This work was supported in part by the Intramural Research Program of the National Institute of Allergy and Infectious Diseases, National Institutes of Health. NK is supported by National Institute of Allergy and Infectious Diseases of the National Institutes of Health under Award Number U01AI136033.

## Supplementary Tables

**Supplementary Table 1**. Calculations of the minimal frequencies of mating competent cells for each L747 x MA37 cross.

**Supplementary Table 2**. Differentially expressed genes for each of the seven cell clusters identified in L747 and MA37 (integrated irradiated and untreated samples) according to *FindMarkers* function in Seurat R package, with the following cutoff values: percentage of cells within a cluster expressing a given gene > 10%, |log2FC| > 0.1 and Wilcoxon Rank Sum test adjusted *p-*value < 0.05). Pct.1: percentage of cells in the cluster expressing a given gene; Pct.2: percentage of cells in all the other clusters expressing a given gene.

**Supplementary Table 3**. List of genes upregulated in both L747 cluster 2 and MA37 cluster 1 with the following threshold cutoffs: percentage of cells within a cluster expressing a given gene > 10%, |log2FC| > 0.1 and Wilcoxon Rank Sum test adjusted *p-*value < 0.05.

**Supplementary Table 4**. List of the primers used in this work.

**Supplementary Table 5**. Summary statistics of whole-genome sequencing read and alignment quality.

**Supplementary Table 6**. Summary statistics of single-cell RNA-sequencing reads and alignment quality.

## Supplementary figures

**Sup. Figure 1**. A-C: Growth curves of L747 after exposure to γ-radiation (A), H_2_O_2_ (B) and MMS (C). D-F: Growth curves of MA37 after exposure to γ-radiation (D), H_2_O_2_ (E) and MMS (F).

**Sup. Figure 2**. Flow cytometry data of a representative double-fluorescent irradiation-facilitated hybrid from the different crosses where each parent expresses either GFP or RFP.

**Sup. Figure 3**. Circos plots showing inferred parental SNP contribution in all *L. braziliensis and L. donovani* sequenced hybrid genomes. Red and green histograms refer to homozygous SNPs specific to the RFP-expressing parental line (R) and the GFP-expressing parental line (G), respectively.

**Sup. Figure 4**. DNA content analysis of irradiation-facilitated hybrids by propidium iodide (PI) staining followed by flow cytometry. Parental strains expressing RFP (in red) or GFP (in green) are considered to have close to diploid genomes (2n) while hybrids generated post-irradiation are close to triploid (3n) or tetraploid (4n). Estimated ploidy was assigned based on both PI staining and whole genome sequencing analysis for hybrids generated from *L. tropica* MoMA (A), *L. donovani* Mondo06 (B), *L. braziliensis* MIR and RIM (C) and *L. infantum*-*L. tropica* IMA (D) crosses. Control *L. tropica* LMA hybrids of different ploidies generated in untreated *in vitro* mating are shown (E).

**Sup. Figure 5:** Description of the different approaches attempted to reduce the ploidy of LMA irradiation-facilitated hybrids (hybrids LMIIirAb4 and IIirBb1).

**Sup. Figure 6**. Transcriptomic heterogeneity in untreated *L. tropica* strains. (A) Variable features selected for the Seurat R package analysis of the scRNA-seq data (3000 genes). (B) UMAP visualization of the untreated *L. tropica* clusters for L747 (left) and MA37 (right) strains. Two biological replicates analyzed are shown for each strain. Cluster denomination is arbitrary and unrelated between the strains. (C-D) Relative single-cell expression of different transcripts (log2FC) associated with procylic promastigotes (C), metacyclic promastigotes (D), stress granules (E) or protein synthesis (F).

**Sup. Figure 7**. Representative flow cytometry analysis used to validate hybrids generated by crossing different combinations of mNG-HAP2^+^ and mNG-HAP2^-^ sorted cells from parental strains expressing either tdTomato or mCherry in the small ribosomal subunit rRNA locus (SSU).

