## Supplementary figures for "Stress conditions promote the mating competency of *Leishmania* promastigotes *in vitro* marked by expression of the ancestral gamete fusogen HAP2"

#### Supplementary Figure 1

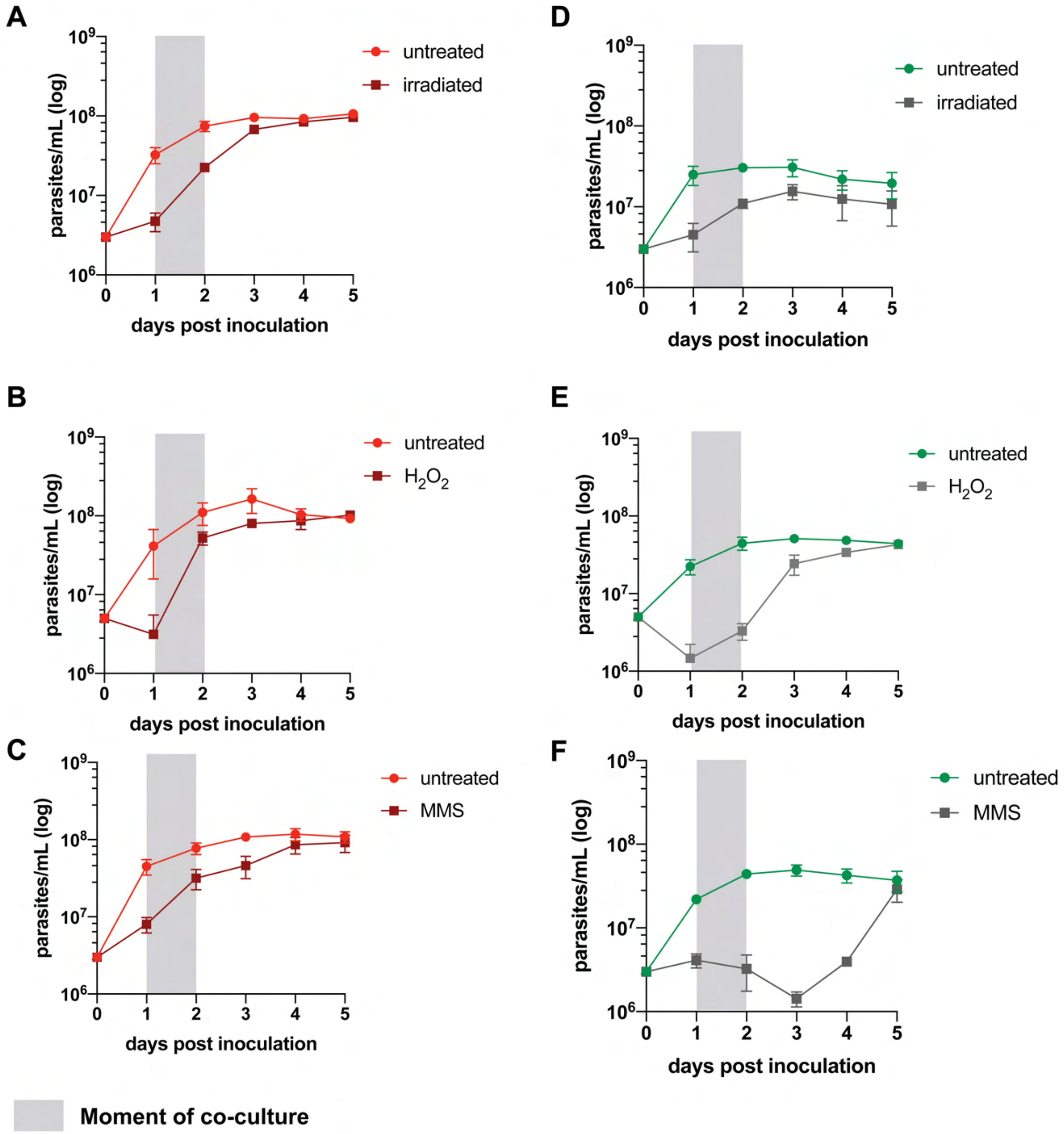

### Supplementary Figure 2

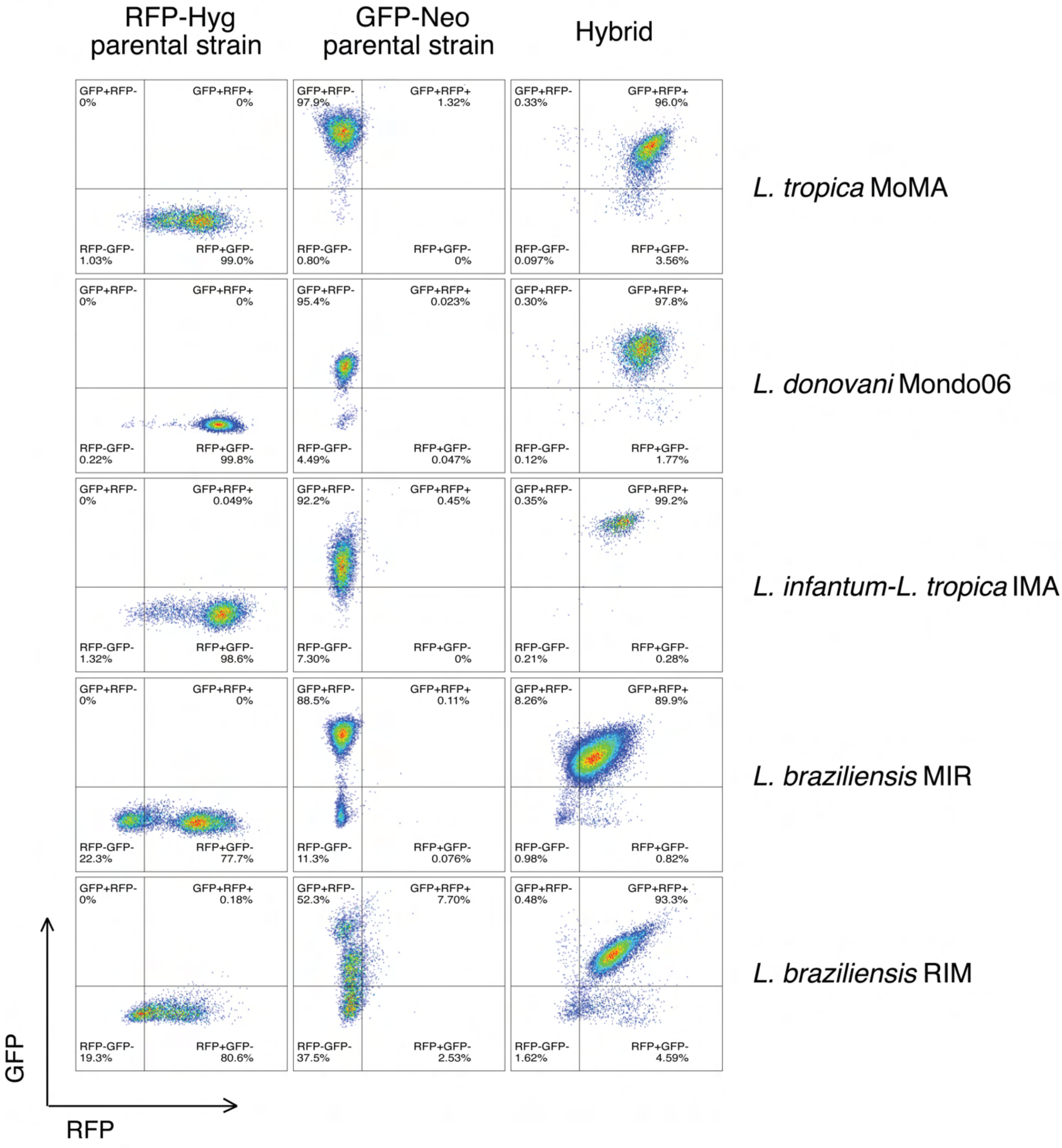

### Supplementary Figure 3

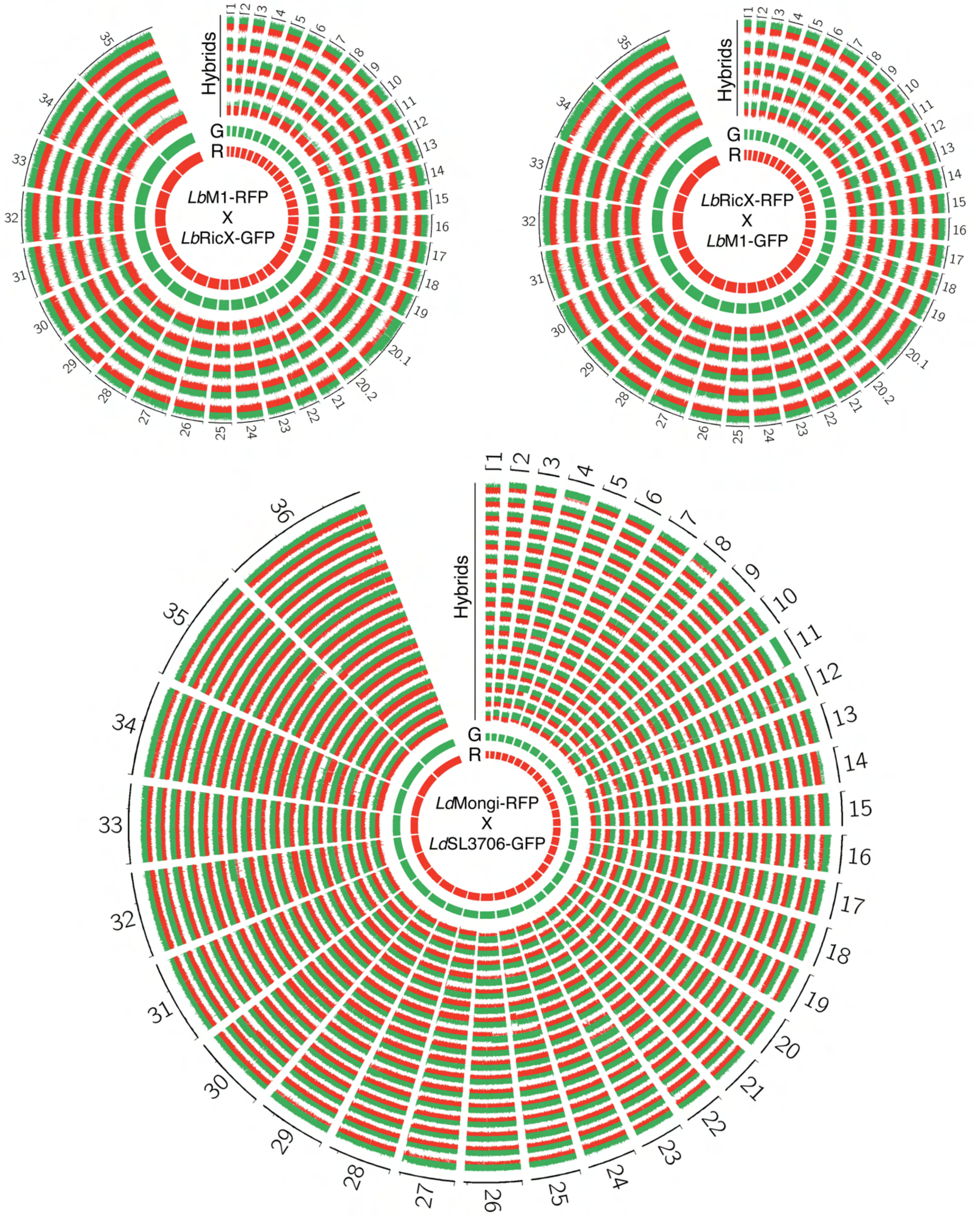

### Supplementary Figure 4

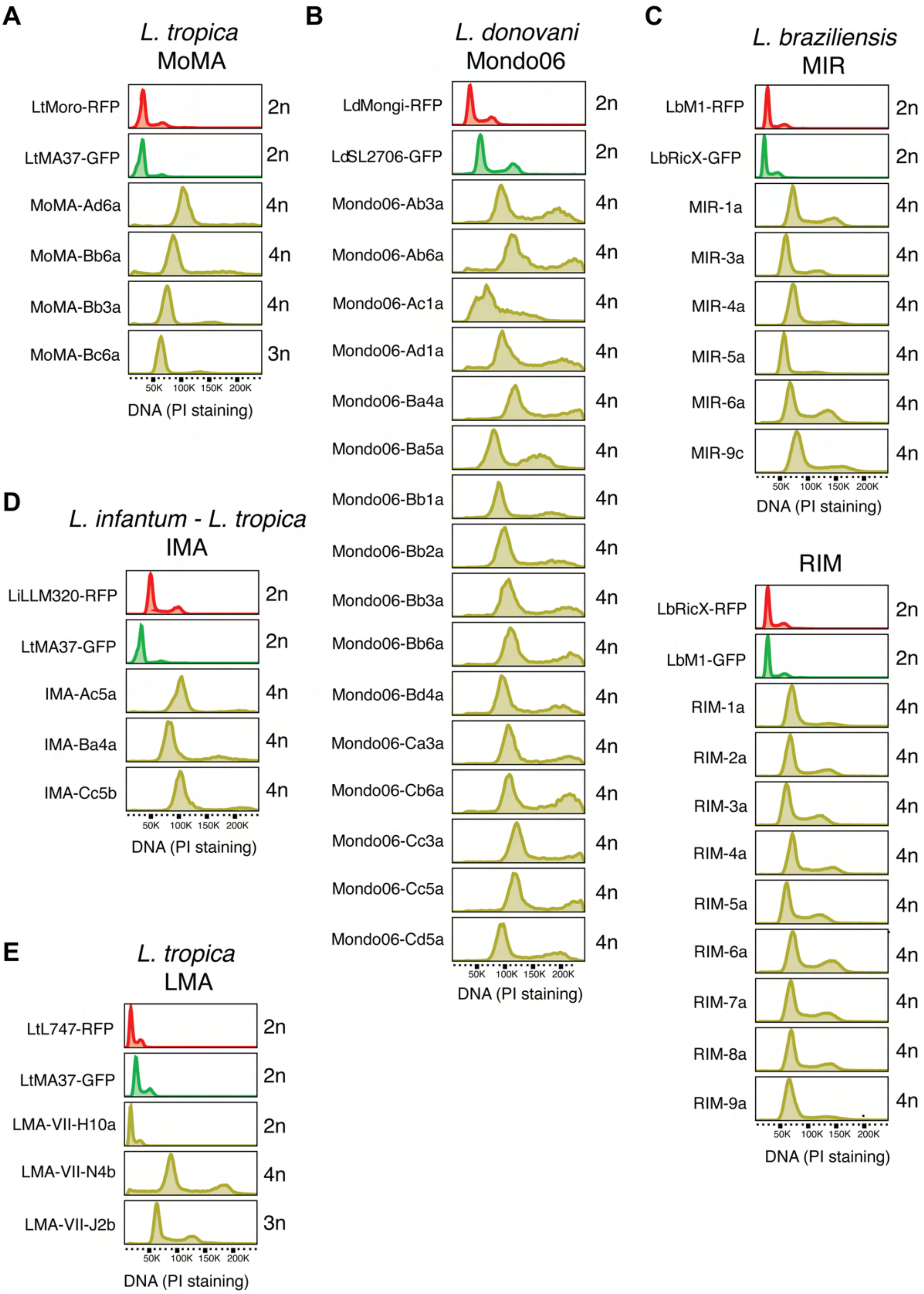

#### Supplementary Figure 5

**Test 1**

**Day 0**

Inoculation in:
- CM199 (control)
- non supplemented M199
- CM199 + $H_2O_2$
- filtered, 3 weeks used CM199

**Day 4**

+ fresh CM199 in all cultures

**Day 7**

DNA content analysis: no visible population of cells with a ploidy <4n in any of the cultures

**Test 2**

**Day 0**

Inoculation in:
- CM199 (control)
- CM199 + $H_2O_2$
- filtered, 3 weeks used CM199
- CM199 + exposure to irradiation

**Day 1**

Subcloning in fresh CM199

**Day 13**

DNA content analysis: no visible population of cells with a ploidy <4n in any of the cultures (x clones tested for each condition)

**Test 3**

**Day 0**

Inoculation in CM199

**Day 1**

Infection L.longipalpis ($4x10^6$/mL of blood)

**Day 9**

- Fly dissection
- Culture of gut extracts

**Day 16**

DNA content analysis: no visible population of cells with a ploidy <4n in any of the cultures (x gut extract cultures)

**Test 4**

**Day 0**

Inoculation in CM199

**Day 1**

Infection L.longipalpis ($4x10^6$/mL of blood)

**Day 9**

- Fly dissection
- 6 flies: immediate subcloning of the parasites extracted from the gut
- 30 flies: culture of gut extract without subcloning

**Day 16-20**

DNA content analysis: no visible population of cells with a ploidy <4n in any of the cultures (x uncloned and x cloned cultures)

### Supplementary Figure 6

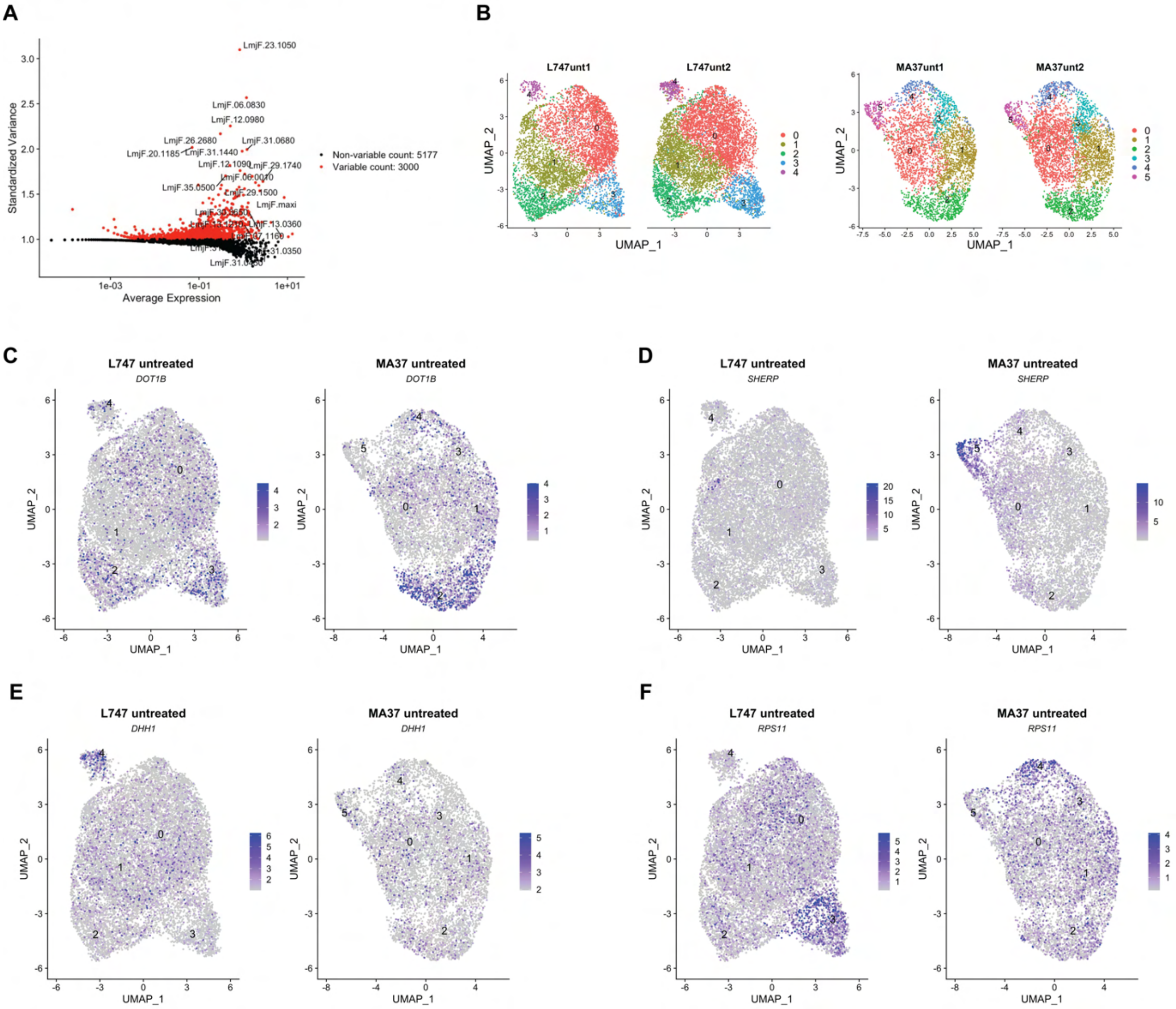

### Supplementary Figure 7

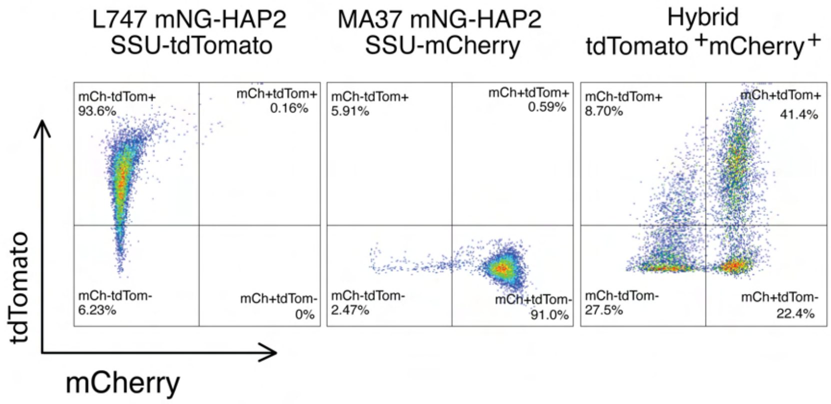


LMA cross
using mNG-HAP2 + or mNG-HAP2⁻
cells sorted from parental strains
